# Defined roles for the *Staphylococcus aureus* POT transporter DtpT in di/tripeptide uptake and glutathione utilisation inside human macrophages

**DOI:** 10.1101/2024.12.02.626324

**Authors:** Imran Khan, Sandy J MacDonald, Sigurbjörn Markusson, Paige J. Kies, Cristina Kraemer-Zimpel, Callum Robson, Joanne L Parker, Simon Newstead, Dave Boucher, Neal D. Hammer, Marjan Van Der Woude, Gavin H Thomas

## Abstract

Peptides available in biological niches inhabited by the human pathogen *Staphylococcus aureus* serve as a rich source of amino acids required for growth and successful host colonisation. Uptake of peptides by *S. aureus* involves at least two transport systems: the di/tripeptide permease DtpT and the oligopeptide ABC transporter Opp3. Here we study the individual and combined functions of DtpT and Opp3 in enabling utilisation of diverse di-/tripeptides via a high-throughput phenotypic screen. We reveal that DtpT is the primary route of uptake for dipeptides, and although many peptides can be utilised via either of the two transport systems, we demonstrate a clear preference for Asp/Glu-containing peptides among DtpT substrates.

To better understand the substrate preferences of DtpT, the protein was purified and reconstituted into proteoliposomes. Active transport of diverse di- and tripeptides was demonstrated, supporting the conclusions of the phenotypic screen. During this *in vitro* analysis, we discovered that DtpT could transport the biologically prevalent tripeptide glutathione (GSH). Bacterial growth assays demonstrate that *dtpT* is essential for GSH utilisation in the absence of the known glutathione transporter, Gis, identifying DtpT as the second GSH uptake system of *S. aureus*. We demonstrate that GSH transport is required by *S. aureus* for complete fitness during *in vitro* macrophage infection experiments. Finally, based on analysis of the DtpT structure and identification of key residues needed for GSH binding and transport, we suggest that GSH transport may be conserved in the DtpT orthologue of *Listeria monocytogenes.* Together, these data reveal important new functions for DtpT in the utilisation of diverse peptides and point toward a novel role for DtpT (and, potentially, other bacterial POT proteins) in glutathione acquisition during intracellular infection.

**Author summary:** The environments where bacterial pathogens thrive are often rich in proteins and their degradation products, including oligopeptides, which can be taken up by the bacterium and used as nutrients. Understanding how this occurs could help us find ways to tackle the growth of these pathogens during infection. Here we examine how the major human pathogen *Staphylococcus aureus* takes up oligopeptides. We demonstrate that a membrane transporter protein, DtpT, is the major route of dipeptide uptake and reveal over 100 new di-/tripeptide targets for this protein. Our findings highlight a defined role for DtpT in the accumulation of Aspartate- and Glutamate-containing peptides, which may serve as relevant nitrogen sources during infection. We also provide the first evidence that DtpT transports the prevalent human metabolite reduced glutathione, and demonstrate that DtpT functions alongside the previously identified Gis glutathione transport system to support intracellular survival of this pathogen inside macrophages. Overall, our findings provide a clear example of how substrate selectivity allows DtpT to fulfil specific biological roles in *S. aureus*, and this functional specialisation may be a common feature of homologous peptide transporters in other bacterial pathogens and across the Tree of Life.

## Introduction

*Staphylococcus aureus* is a Gram-positive human bacterial pathogen associated with a broad range of different disease manifestations including skin/soft tissue infections, systemic bacteraemia, endocarditis and other invasive disease states associated with significant mortality (Tong et al., 2015). In order to meet their nutritional requirements during infection, bacterial pathogens such as *S. aureus* possess diverse transport systems which allow for the selective uptake of nutrients present within the host. Understanding the intrinsic factors contributing to bacterial fitness, colonisation and infection in relevant infection environments is essential for informing the design of novel therapeutic approaches. While the importance of some individual transporters in successful colonisation is well documented – particularly those implicated in the acquisition of key carbon sources, amino acids and metals (Adolf et al., 2023; Handke et al., 2018; Lehman et al., 2023; Lensmire et al., 2020) – the roles of many transport systems and their contributions toward infection remain understudied.

Peptides serve as an abundant source of amino acids, and hence can provide carbon, nitrogen and sulphur. As such, most bacterial pathogens possess one or more routes of uptake for these molecules. Most bacterial peptide transporters studied to-date are members of the ATP-Binding Cassette (ABC) family of importers or secondary active transporters belonging to the Major Facilitator Superfamily (MFS), with the latter being further divided into a small number of functionally distinct groups including the proton-dependant oligopeptide transporters, or POTs (Garai et al., 2017). ABC peptide importers are unique to prokaryotes and are capable of recognising and transporting diverse oligopeptides with high affinity, with substrates ranging from di-/tripeptides up to 35-mers (Detmers et al., 2000; Doeven et al., 2004; Hughes et al., 2022; Smith et al., 1999). In addition to their canonical role in nutrient acquisition, there are examples of these transporters contributing to diverse physiological functions including cell-wall turnover (Hughes et al., 2022; Maqbool et al., 2011) and antimicrobial peptide resistance (Ackroyd et al., 2024; Wang et al., 2016), as well as acting as sensory units for intercellular and environmental signalling networks (Slamti & Lereclus, 2019). POTs, by comparison, are found across all domains of life and exclusively transport di- and tripeptides or chemically related molecules of a similar size (Newstead, 2017; Kotov et al., 2023; Parker et al., 2024). Despite their ubiquity, the importance of bacterial POTs during infection is poorly characterised and little is known regarding their contribution to bacterial physiology outside of their general role in nutrient uptake.

To date, there are two known peptide transport systems which are widely distributed in Staphylococci: the ABC importer-type system Opp3_ABCDF_ (or Opp3) and the di-/tri-peptide POT transporter DtpT (Yu et al., 2014). In the single elegant study comparing the roles of these two systems in *S. aureus*, Opp3 was demonstrated to be required for the transport of oligopeptides from 3 to 9 residues in length, while only DtpT was thought to be responsible for dipeptide uptake (Hiron et al., 2007). *S. aureus* also possesses an ABC importer system which is specific for the biologically prevalent tripeptide glutathione (Gis_ABCD_, or Gis) (Lensmire et al., 2023). In addition to the aforementioned systems, *S. aureus* strains can possess up to four additional Opp3-homologues, of which Opp1 and Opp2 are likely metal rather than peptide transporters (Hebbeln & Eitinger, 2004; Remy et al., 2013). However, Opp4 plus one additional Opp system (ACME Opp) encoded as part of the widely disseminated arginine catabolic mobile element (Yu et al., 2014) are mostly uncharacterised and could function as additional peptide transporters.

The importance of peptide utilisation in *S. aureus* infection biology is a subject of ongoing investigation. *S. aureus* secretes an arsenal of extracellular proteases with diverse functions in host immune modulation and virulence, and these enzymes contribute to the generation of peptides within the local environment which are likely to serve as a nutrient source for the bacterium (Coulter et al., 1998; Shaw et al., 2004). Consistent with this, there is evidence that *S. aureus* senses protein-rich environments via Opp3 and responds to this stimulus via upregulation of extracellular proteases (Borezée-Durant et al., 2009; Lehman et al., 2019). Recent work has demonstrated how the *S. aureus* protease aureolysin is capable of contributing to the degradation of collagen and highlights how growth on collagen-degradation products is decreased in an Opp3-deficient mutant (Lehman et al., 2019). These observations point toward a model in which host- and bacterial-derived peptides support the growth of *S. aureus* in an Opp3-dependent manner. Conversely, loss of DtpT – but seemingly not Opp3 – leads to attenuation in several models of invasive *S. aureus* infection including intravenous infection in mice and an endocarditis model in rabbits (Coulter et al., 1998). In spite of this, the functional characterisation of DtpT to date is limited, and the molecular basis of its role in bacterial survival during infection is unknown.

Recent work has aimed to understand the molecular basis of substrate promiscuity observed in POTs and delineate the mechanistic basis of proton-coupled peptide transport in these proteins (Kotov et al., 2023; Lichtinger et al., 2024; Parker et al., 2014). However, little is known regarding the substrate preferences of these transporters and how this relates to cellular lifestyle/environment. Here, we aim to address this gap in our knowledge. We apply Phenotype Microarrays (BIOLOG) to compare the utilisation of varied di- and tripeptides by wild-type *S. aureus* JE2 against strains with disruptions in the known peptide transporters. This approach provides novel insights into the diversity of peptides utilised via DtpT and points toward distinct amino acid preferences among DtpT substrates. Further cell-based and biochemical assays serve to both support these findings and further interrogate the specificity of the system. Finally, we extend these assays to demonstrate how DtpT also serves as a transporter for the biologically prevalent metabolite reduced glutathione (GSH) and demonstrate how glutathione transport mediated by DtpT and Gis is a determinant of bacterial survival inside human macrophages.

## Results

### Phenotypic screening defines the substrate landscapes of DtpT and Opp3A

In order to investigate the roles of the two known peptide transporters in *S. aureus*, we generated an isogenic series of strains in the USA300-derivative strain JE2 with disruptions of *dtpT* and *opp3A,* as well as a double-mutant strain. With these strains in hand, we exploited the power of phenotype microarrays (PMs) as a simple high-throughput approach for comparing the abilities of bacterial strains to catabolise diverse substrates, including di- and tripeptides (Biolog PM Plates 6–8) (Bochner et al., 2001; Pletzer et al., 2014). Collectively, these plates allowed us to compare utilisation for 282 different peptides, comprising: 246 standard dipeptides (of a possible 400), 1 hydroxyproline-containing dipeptide, 7 β-bonded dipeptides, 2 γ-bonded dipeptides, 12 D-amino acid-containing dipeptides and 14 tripeptides. In this assay, a given peptide is supplied as the primary nitrogen source, and cellular metabolism is measured via reduction of a redox-sensitive colourimetric dye, quantified over time to produce a signal curve for each strain/peptide combination. A positive metabolic signal is indicative of peptide utilisation enabled by a valid route of uptake and downstream catabolic pathways, while intermediate/weak signals are likely to result from poor uptake (or a complete lack thereof) or incomplete catabolism following internalisation.

Based on the signal curves produced from our PM analysis, we could observe a series of distinct utilisation patterns across our four test strains (Supplemental data SD1-4). To identify commonalities in the data, we subjected the raw signal curves for each peptide to unbiased cluster analysis and dimensional reduction using Uniform Manifold Approximation and Projection (UMAP) (Figure 1). These analytical methods provided complementary outputs, with peptides belonging to a common cluster demonstrating close spatial organisation in the final UMAP plot (Figure 1A-C).

**Figure 1.**
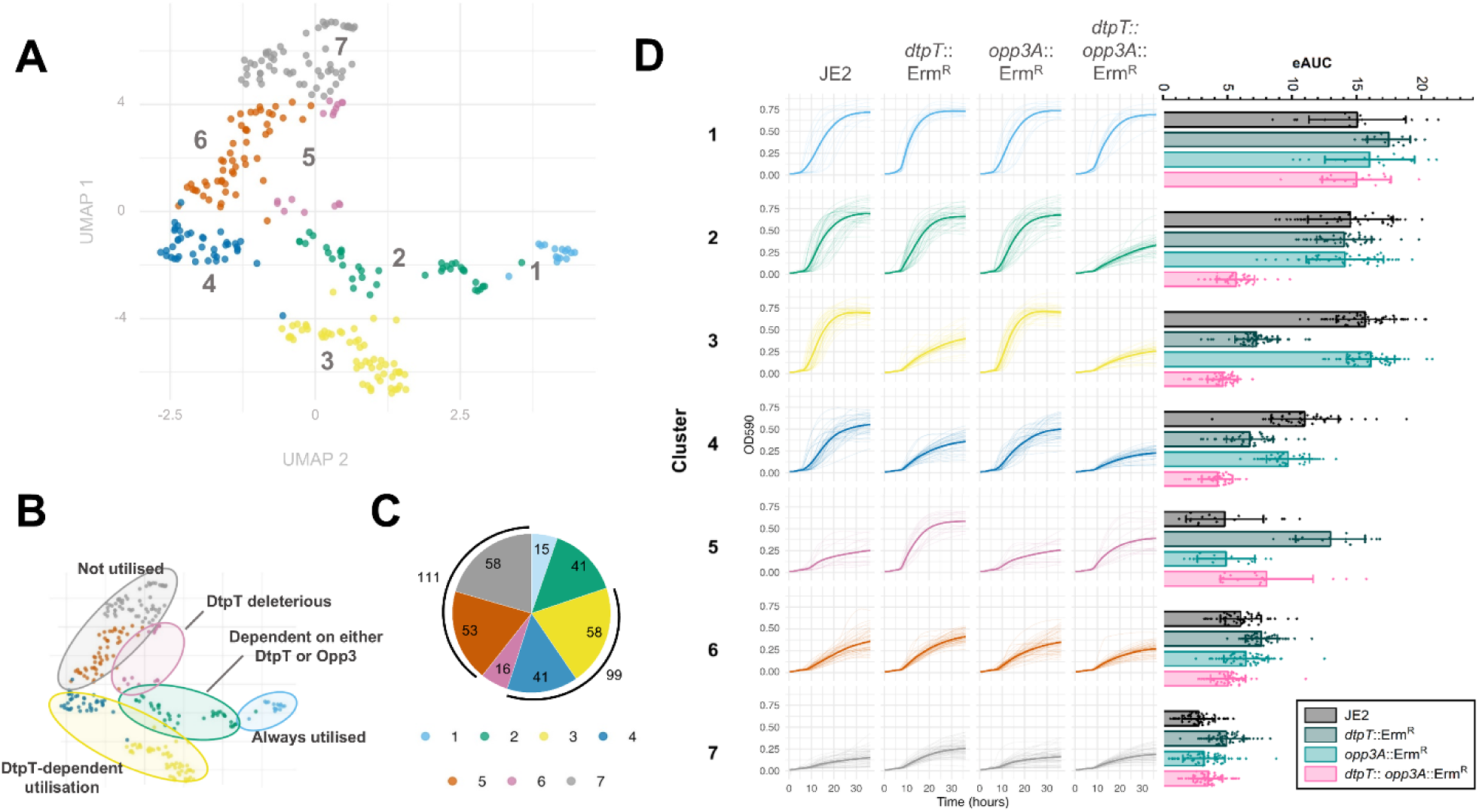
UMAP and cluster analysis reveal differential peptide utilisation patterns across *dtpT* and *opp3A* mutant strains. (**A**) PM signal curves were subjected to UMAP and k-means cluster analysis (k = 7). Peptides belonging to the same cluster are grouped by colour on the UMAP plot, as labelled. (**B**) Regions of the UMAP plot corresponding to distinct phenotypic patterns (determined by clustering) are highlighted. (**C**) Pie-chart representation of the number of peptides in each of the seven clusters identified here. In each case, colours correspond to the seven clusters identified in this analysis, as depicted by the key. (**D**) (**Left**) Cluster analysis reveals seven strain-specific peptide utilisation patterns. All signal curves for each cluster are overlayed and separated by strain. Mean signal curves for each strain are indicated by the bold line. (**Right**) Empirical area under the curve (eAUC) was calculated for all signal curves. Bars show the mean eAUC for each strain within a given cluster. Error bars indicate standard deviation in each case.

Visual confirmation of the clusters suggested that each encompasses a distinct pattern which reflects differences in the routes of peptide utilisation (Figure 1B/D). Cluster 1 (n = 15) contains peptides for which a strong metabolic signal was observed for JE2 and all three mutant strains, suggesting – rather intriguingly – that neither of the two known peptide transporters of *S. aureus* is required for their uptake. In contrast, utilisation of peptides in cluster 2 (n = 41) was maintained in single mutant strains lacking either *dtpT* or *opp3A* but abolished in the double-mutant strain, suggesting that these peptides can be transported by either of our two systems. Clusters 3 and 4 (n = 99) both present patterns wherein utilisation is abolished for the *dtpT* mutant and the double mutant, but not the *opp3* mutant, suggesting DtpT serves as the primary route of uptake for these peptides. Collectively, these two clusters account for more than one-third of the tested peptides, with those in cluster 3 (n = 58) demonstrating a more pronounced phenotype (that is, a greater difference in signal between “positive” and “negative” utilisation) when compared to those in cluster 4 (n = 41). Such a distinction likely results from either a higher rate of internalisation by DtpT or more productive downstream catabolism following uptake for cluster 3 peptides.

The three remaining clusters contain peptides for which a weak metabolic signal was observed in the wild-type JE2 strain. Cluster 5 (n = 16) encompasses peptides for which, quite surprisingly, metabolism is permitted only in mutant strains lacking *dtpT*. While the exact mechanistic basis for this pattern is unclear, we postulate that the DtpT-mediated internalisation of these peptides may be cytotoxic under the conditions tested. However, a strong signal in the *dtpT* mutant strain suggests an alternate route of internalisation for which toxicity is not observed, perhaps due to a lower overall flux which prevents toxic over-accumulation of the peptide. The remaining peptides each gave a weak/intermediate signal or were not utilised at all or across the four strains tested here, as observed in cluster 6 (n = 58) and cluster 7 (n = 53) respectively. These phenotypic patterns suggest that the supplied peptide alone cannot be effectively utilised for cellular metabolism. We do note that the mean eAUC values recorded for cluster 6 are slightly higher than those in cluster 7 across the four strains, perhaps indicating that some peptides in cluster 6 can be utilised to produce an intermediate metabolic signal.

Notably, our analysis did not identify a cluster (nor even any individual peptides) for which the patterns of metabolism suggested that utilisation was solely dependent on Opp3, which is perhaps expected given that this system is primarily implicated in the utilisation of larger oligopeptides (n ≥ 3) (Hiron et al., 2007; Lehman et al., 2019). Taken together, our findings implicate DtpT in the utilisation of at least 140 of the peptides assessed here, with 41 of these being recognised by both peptide transport systems.

### Cluster analysis demonstrates amino acid preference among putative DtpT substrates

Next, we aimed to determine whether specific amino acids were enriched within the peptides of each cluster. Given the limited panel of tripeptides assessed, we focussed our analysis on the 246 standard dipeptides present in the assay. First, we plotted a grid to identify potential cluster-specific trends in regard to the N- and C-terminal amino acid composition of dipeptides included in the phenotype microarray analysis (Figure 2). Then, we examined any statistically meaningful relationships by applying a Fisher exact test (Figure 3A-D). The strongest association was identified between glutamine-containing peptides and cluster 1, with the vast majority (9/13) of Gln-containing peptides assessed here belonging to this cluster (Figure 3A). Such an observation strongly suggests that an alternate route of uptake exists for Gln-containing peptides in *S. aureus,* which is independent of both DtpT and Opp3. Another clear relationship was observed between tyrosine-containing dipeptides and cluster 7 (Figure 3C), suggesting that tyrosine-containing peptides are either poorly transported or metabolised by *S. aureus* under the conditions tested here. Tantalisingly, we observe that aspartate and glutamate-containing peptides are both strongly enriched within the DtpT-specific cluster 3 (Figure 3D). This preference is seemingly independent of amino acid arrangement, with comparable frequencies for DtpT-specific utilisation in the N- or C-terminal positions (Figure 2). Our findings provide compelling evidence that Asp- and Glu-containing peptides are preferred transport targets for DtpT, in-turn suggesting that utilisation of these peptides is a distinct function of the transporter.

**Figure 2.**
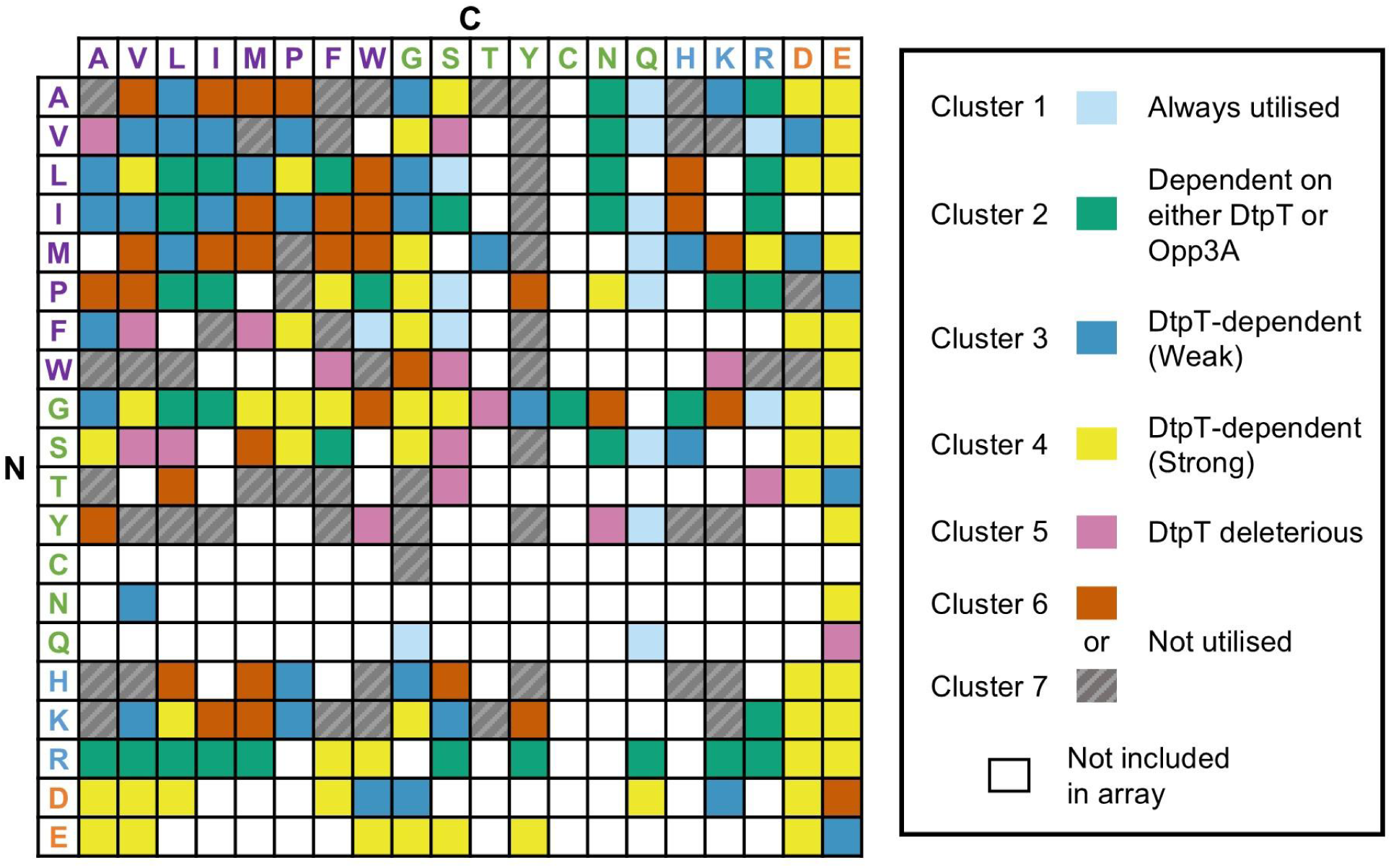
Phenotype microarrays reveal differential routes of uptake for diverse dipeptides in *S. aureus.* Grid representation of all possible dipeptide structures arranged with N-terminal amino acids on the Y axis and C-terminal amino acids on the X axis. Amino acids are displayed as single-letter symbols and colour-coded according to their chemical properties (purple = hydrophobic/non-polar, green = polar/unique, blue = basic, orange = acidic). Coloured cells correspond to peptides included in the PM analysis and their corresponding utilisation patterns, as indicated by the key.

**Figure 3.**
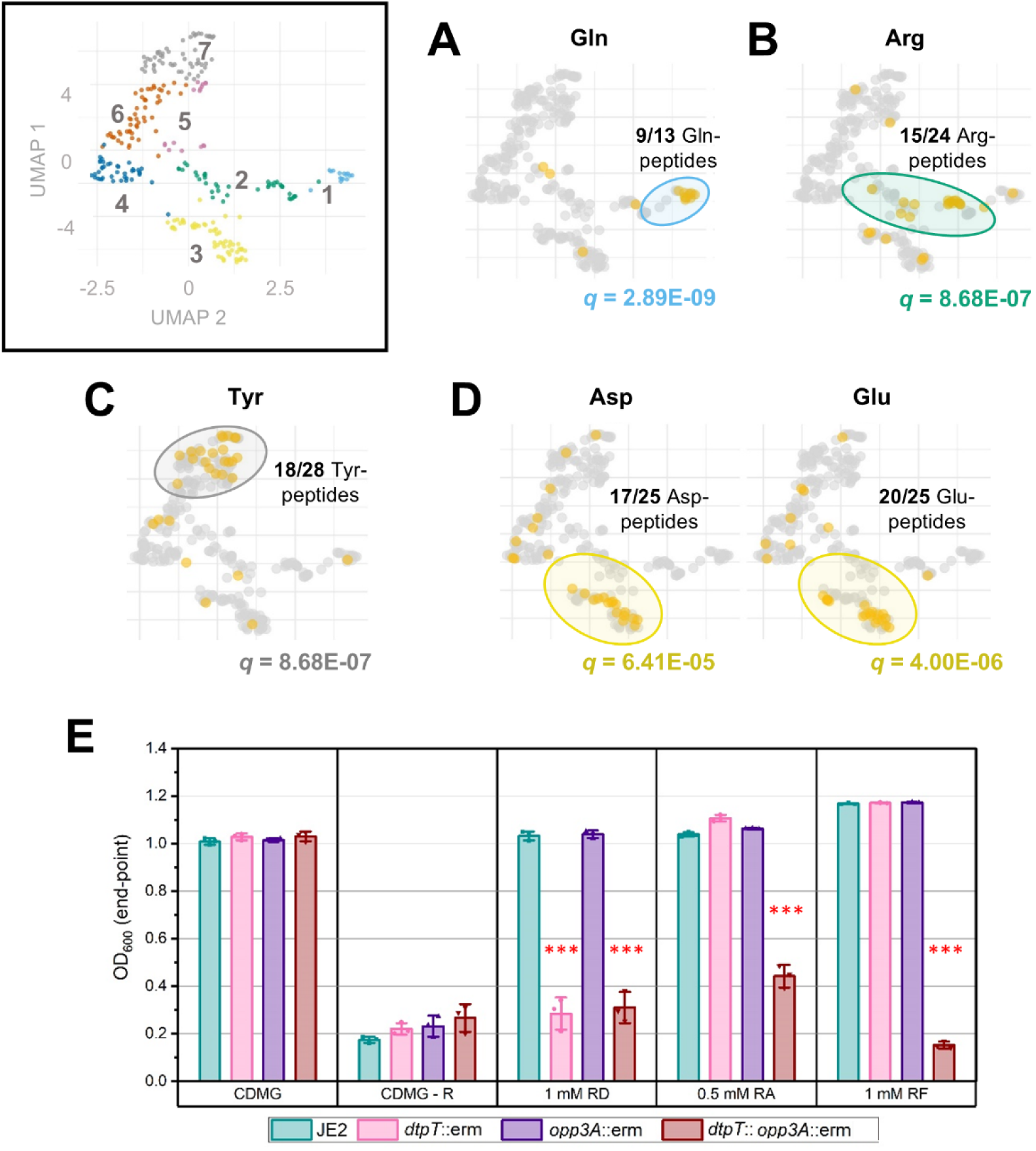
Specific amino acids are enriched in peptides belonging to a common utilisation pattern. (**A - D**) Enriched amino acids within each cluster were identified via a Fisher exact test with FDR q-value correction. Plots show the position of peptides containing strongly enriched amino acids (*q* ≤ 1E-03) overlayed on the original UMAP plot (as labelled). The value of *q* is given in each case. (**E**) Growth of *S. aureus* strain JE2 and mutant derivatives in CDMG – R supplemented with three dipeptides, as labelled. Arg-Asp (RD) and Arg-Phe (RF) are each cluster 3 peptides, while Arg-Ala (RA) belongs to cluster 2. Bars show the mean end-point OD_600_ value after 24 hours of growth ± standard deviation. End-point OD is reflective of the maximal cell density observed over this period. *** p < 0.001; unpaired t-test (vs JE2).

In addition, we found that arginine-containing peptides were highly enriched in cluster 2 (Figure 3B), implicating both peptide transport systems in the utilisation of these peptides. Arginine auxotrophy in *S. aureus* USA300 was previously demonstrated using chemically defined medium containing glucose (CDMG) and the same medium lacking arginine (CDMG-R) (Jeong et al., 2022; Reslane et al., 2022). Here, we exploit this auxotrophy to verify the DtpT/Opp3A-dependence patterns of three arginine-containing peptides as observed in our phenotypic screen. Specifically, we supplemented CDMG-R with two cluster 3 peptides (Arg-Asp and Arg-Phe) and one cluster 2 peptide (Arg-Ala) and separately inoculated these media with JE2 and the three mutant-derivative strains. As expected, robust growth for all strains was observed in CDMG, while none of the strains were able to grow in CDMG-R (Figure 3E; Figure S1A). Supplementing with Arg-Asp (cluster 3) restored growth in a DtpT-dependent manner, while Arg-Ala (cluster 2) restored growth in all but the double mutant strain (Figure 3E; Figure S1B). These patterns support the findings of our phenotypic screen. Unexpectedly, supplementing with Arg-Phe (cluster 3) restored growth in all but the double mutant strain similarly to Arg-Ala (cluster 2). Such deviation is likely indicative of differences in the sensitivity of the two approaches utilised here; that is, Opp3-mediated transport of Arg-Phe is sufficient to relieve Arg auxotrophy when supplied at 1 mM, but too limited to support metabolism as a sole nitrogen source under PM conditions. Finally, to further verify the role of *dtpT* in the utilisation of these three peptides, we cloned the gene into a low-copy number vector and were able to observe genetic complementation of growth in each case (Figure S1C). Hence, we confirm that DtpT serves as a route of entry for cluster 2 and cluster 3 peptides, supporting the findings of our PM assays.

### DtpT transports peptides with diverse chemical properties

Peptide utilisation (as quantified by phenotype microarrays) produces a composite phenotype representing the combined consequences of peptide internalisation and catabolism, both of which are required to drive cellular metabolism and produce a positive signal. As a result, peptides with limited uptake may produce a positive PM signal if efficiently catabolised while other peptides may be rapidly internalised (by DtpT or otherwise) without producing a positive PM signal due to incomplete catabolic pathways.

To compare DtpT-mediated transport of peptides directly, we expressed and purified *S. aureus* DtpT recombinantly from *E. coli* cells (Figure S2) and reconstituted the pure protein into liposomes (Figure S3A-B), using methods previously applied to study the *Staphylococcus hominis* DtpT orthologue PepT_Sh_ (Minhas et al., 2018). Transport of di-/tripeptide substrates was then assessed by utilising the pH-sensitive dye pyranine, as described previously (Parker et al., 2014). Briefly, peptide transport was assayed by monitoring the acidification of the liposome lumen and quantified over time for each peptide after induction of a negative-inside membrane potential (ΔΨ) (Figure 4A). To validate this methodology, we confirmed that peptide transport is dependent on the presence of both a valid substrate and ΔΨ, as evidenced by the lack of strong acidification when either of these components is absent (Figure S3C) and were then able to measure transport for a range of selected di- and tripeptides (Figure 3B-C).

**Figure 4.**
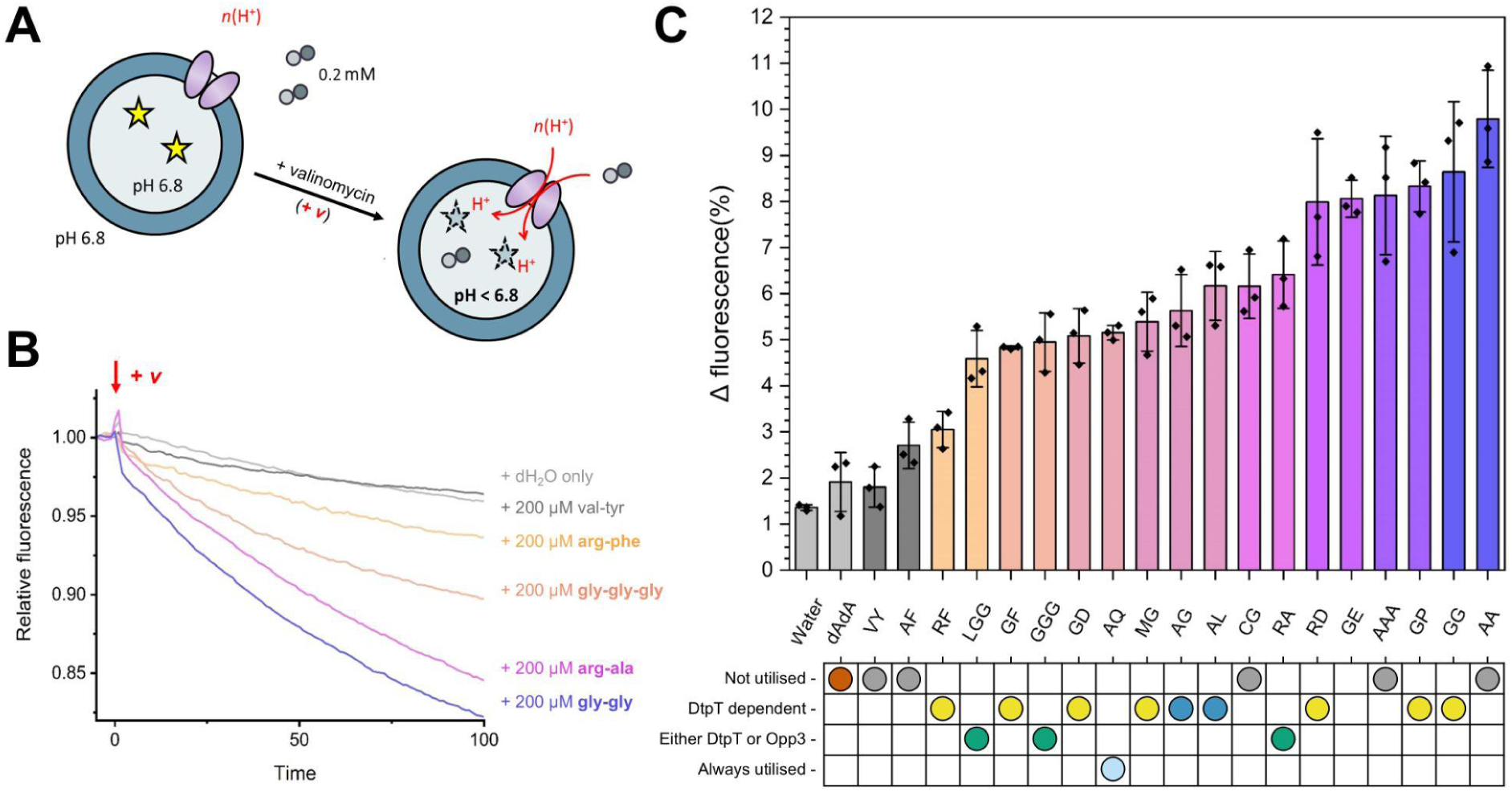
DtpT transports structurally diverse di-/tripeptides. (**A**) DtpT was reconstituted into liposomes containing pyranine. Upon induction of negative-inside ΔΨ, DtpT transports peptides from the external environment into the liposome lumen. Peptide transport is proton-coupled, leading to acidification of the lumen and a decrease in the pyranine fluorescence (ex. 460 / emm. 510). (**B**) Exemplary transport data for 5 peptides normalised immediately before the addition of valinomycin (red arrow) for ease of comparison. Curves show the mean of three replicates. (**C**) Transport of 20 peptides was compared by quantifying the change in pyranine fluorescence over the first 30 seconds of the assay. Bars indicate the mean of three replicates ± standard deviation. Coloured bars indicate a change in fluorescence which differs significantly from the no peptide (“Water”) control, as determined by a one-way ANOVA with Fisher LSD test (p ≤ 0.05). Peptides are labelled according to their single-letter amino acid code (dAdA = d-ala-d-ala). Corresponding phenotypic patterns identified by PM analysis for each peptide are shown below and coloured (as established in Figure 1).

By assessing transport of 20 peptides with diverse chemical properties, we were able to confirm DtpT-mediated transport for 12 of the putative substrates identified by our phenotype microarrays (Figure 4C). Notably, we confirm DtpT-mediated transport of the three Asp/Glu-containing dipeptides tested here, supporting the previous observation that peptides in this category are preferred DtpT substrates. We also confirmed the transport of Arg-Phe, Arg-Ala and Arg-Asp by DtpT, as already demonstrated in our Arg-auxotrophy experiments. Overall, there is good agreement between our liposome-based transport assays and whole-cell peptide utilisation methods (Figure 4C). We do note that some peptides are transported by DtpT which did not sustain metabolism of *S. aureus* in the phenotype microarray assay, namely; Ala-Ala, Ala-Ala-Ala and Cys-Gly. Such an observation suggests that internalisation is not the limiting factor in utilisation of these peptides.

Additionally, our transport assays shed new light on the molecular factors contributing toward substrate preference in this transporter. For instance, our data demonstrate how many favourable DtpT substrates incorporate acidic amino acids (Arg-Asp and Gly-Glu) or amino acids with small, uncharged side chains (Ala-Ala, Gly-Gly and Gly-Pro) (Figure 4C). In contrast, we demonstrate that dipeptides containing aromatic residues in position 2 generally demonstrate poorer transport when compared to those with less bulky sidechains in the equivalent position. Poor internalisation of these substrates (e.g. Val-Tyr and Ala-Phe) by DtpT is consistent with their lack of utilisation in PM assays (Figure 4C). Overall, our transport assays demonstrate that DtpT is highly promiscuous and capable of transporting substrates with a range of chemical structures and properties, which is similarly noted for other POT transporters studied previously (Kotov et al., 2023; Parker et al., 2014).

### DtpT is a functional glutathione permease

Reduced glutathione (GSH) is a biologically important tripeptide metabolite with diverse roles in many organisms, primarily serving as an antioxidant in eukaryotes and some prokaryotes (Lushchak, 2012). GSH consists of a Cys-Gly dipeptide conjugated to glutamate via a γ-linked peptide bond between the glutamate side chain and amino terminus of the cysteine (Figure 5A). Given that DtpT recognises and transports Cys-Gly (Figure 4C), we predicted that DtpT may also transport GSH and that this activity may be relevant in biological environments where glutathione is enriched. To test this, we measured DtpT transport activity with either GSH or oxidised glutathione (GSSG) (Figure 5B). We demonstrate that GSH is transported by DtpT, with acidification of the liposome lumen over the first 30 seconds comparable to that observed for Cys-Gly. We did not observe transport of GSSG in this assay, likely due to the larger structure of GSSG compared to GSH.

**Figure 5.**
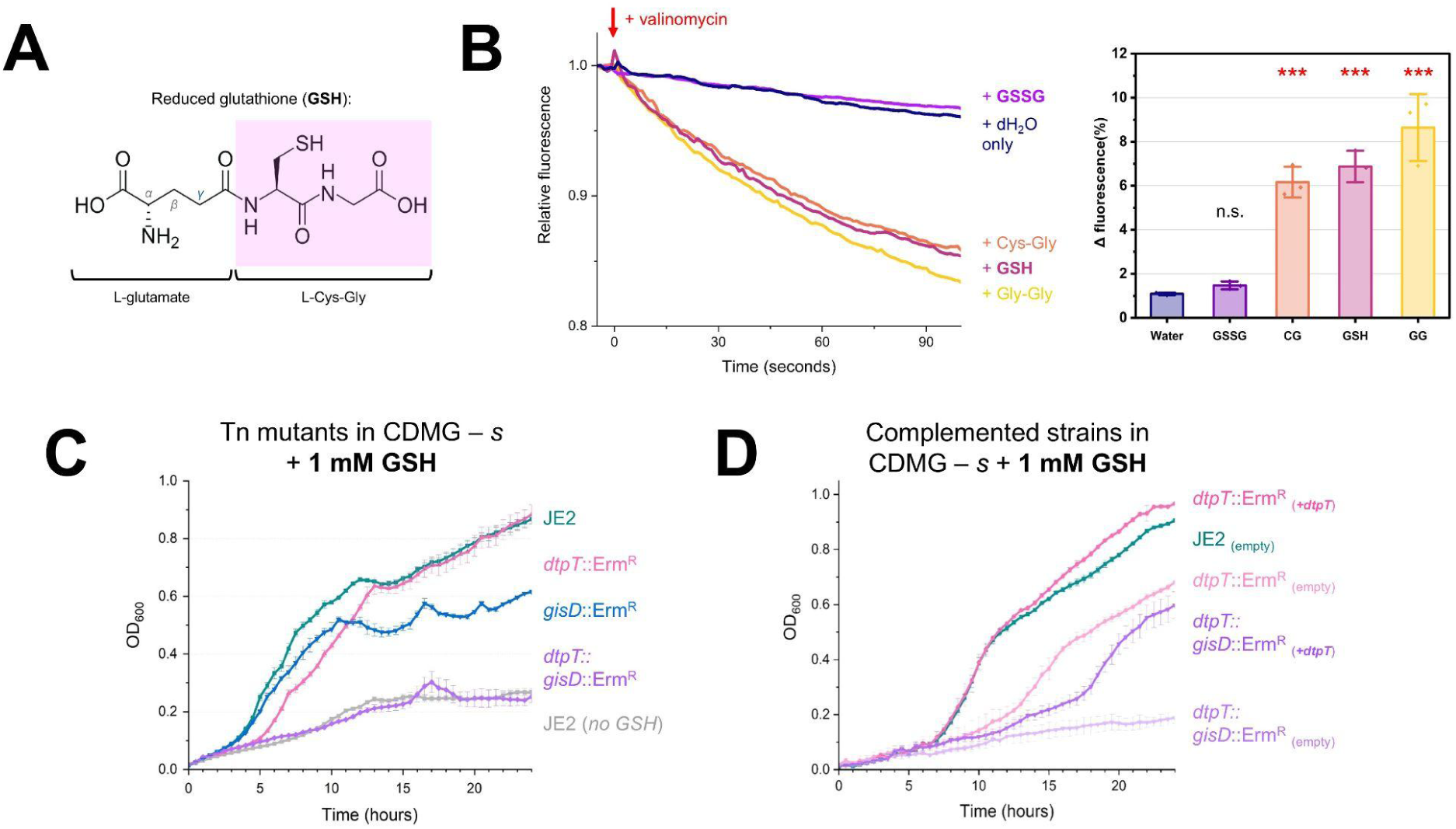
DtpT transports reduced glutathione (GSH). (**A**) Chemical structure of GSH. (**B**) (**Left**) Transport of GSH and GSSG was assessed via pyranine transport assays. Transport assay curves are normalised immediately before the addition of valinomycin (red arrow) for ease of comparison. Curves show the mean of three replicates. (**Right**) summarised transport data. Bars show the mean of three replicates ± standard deviation. ***p<0.001; Unpaired t-test (vs Water). (**C - D**) Growth of *S. aureus* strain JE2 and mutant derivatives was assessed in CDMG – *s* supplemented with 1 mM GSH. In each case, strains were grown over 24 hours and OD_600_ was measured every 30 minutes for three biological replicates. Curves indicate the mean values ± standard deviation. (**C**) Growth of single-mutants of *dtpT* and *gisD* as well as a *dtpT gisD* double-mutant strain in CDMG – *s* ± 1 mM GSH. (**D**) *In-trans* expression of DtpT restores growth of the *dtpT* single mutant to wild-type levels and partially restores growth in the *dtpT gisD* double mutant.

The Gis system is an ABC-type importer which has recently been characterised as a route of glutathione acquisition in *S. aureus* and is the only route of uptake for both GSH and GSSG identified in this bacterium to date (Lensmire et al., 2023). However, under conditions where GSH utilisation is essential, the growth of a *gis*-deficient mutant strain is restored to near wild-type levels when GSH is supplied at concentrations greater than 0.5 mM (Lensmire et al., 2023). This implies the existence of at least one additional route of GSH uptake. To determine whether DtpT fulfils this role, isogenic mutants of *dtpT* and *gisD,* as well as a double mutant, were grown under conditions where glutathione was supplied as the sole sulphur source. At 50 µM GSH, we found that the growth of *S. aureus* is limited and solely dependent on the Gis system (Figure S4A). However, both single mutant strains demonstrated growth defects relative to the wild type at 1 mM GSH, while growth was abolished for the double mutant under this condition (Figure 5C). In comparison, we found that growth in the presence of GSSG was solely dependent on Gis (Figure S4B). We also found that *in trans* complementation with *dtpT* was sufficient to fully restore growth of the *dtpT* single-mutant and partially restore growth in the double mutant strain in the presence of 1 mM GSH (Figure 5D). Finally, given that GSH oxidises spontaneously over time in aerated environments, we repeated the growth experiment with 1 mM GSH under anaerobic conditions to limit changes in the oxidation state of GSH during growth (Figure S4C). Again, we observed phenotypes which were consistent with those observed during aerobic growth. Taken together, these observations demonstrate that both Gis and DtpT contribute to GSH uptake at millimolar concentrations.

### Glutathione transport contributes to bacterial survival inside macrophages

Although *S. aureus* was historically considered an extracellular pathogen, recent work has demonstrated that this bacterium survives inside phagocytic cells, highlighting how its intracellular lifestyle may contribute to dissemination and immune evasion during systemic infection (Flannagan et al., 2016; Kubica et al., 2008). To date, there is a poor understanding of the molecular factors governing fitness in these intracellular environments, particularly regarding nutrient availability and acquisition. Cytosolic concentrations of GSH in human cells range from 1 to 10 mM depending on the cell type, making it the most abundant intracellular non-protein thiol (Cui et al., 2023; Meister, 1988). Based on this, we hypothesised that glutathione utilisation may contribute toward the fitness of *S. aureus* in intracellular environments.

Differentiated THP1 macrophage-like cells and human monocyte-derived macrophages (hMDMs) were each infected with either JE2 or mutant derivatives of *dtpT* and/or *gisD*, and bacterial survival was quantified over the first 8 hours post-infection (Figure 6). To account for any changes in fitness resulting from the inclusion of the *bursa aurealis* transposon insertion sequence, we also assessed the survival of a mutant strain carrying the transposon in a presumed phenotypically null locus corresponding to a predicted IS3-family transposase gene (SAUSA300_0060). In THP-1 cells, survival of both the *dtpT* and *gisD* single-mutant strains was reduced slightly on-average at 8 hours but did not differ significantly when compared to the wild-type. In comparison, survival of the *dtpT gisD* double mutant strain was significantly lower at this time-point, being reduced more than two-fold relative to the wild-type (Figure 6A). Similarly, survival of both the *gisD* single-mutant and *dtpT gisD* double-mutant strains was significantly reduced at this time-point in hMDMs derived from eight independent donors (Figure 6B). Overall, these data suggest that GSH transport is required for complete fitness of *S. aureus* during macrophage infection, though the presence of either *dtpT* or *gisD* is seemingly sufficient to restore survival to near wild type levels.

**Figure 6.**
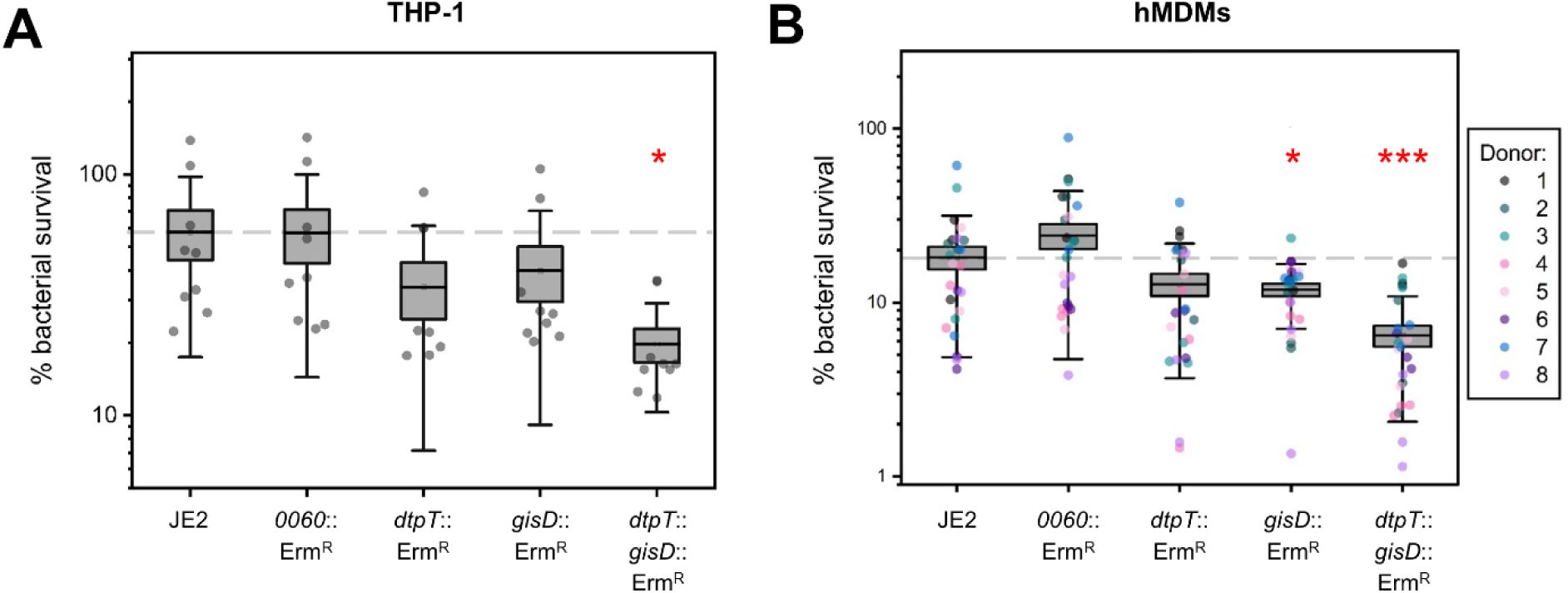
A glutathione transport-null double-mutant demonstrates reduced survival inside macrophages. (**A - B**) Bacterial % survival of JE2 and mutant derivatives after 8 hours post-infection in THP1 macrophage-like cells (**A**; *n* = 9) and hMDM cells (**B**; *n* = 8 donors, each infected in biological triplicate). In each case, survival is calculated as a percentage of the total bacterial uptake. A dashed line indicates the mean % survival of the wild-type JE2. *p<0.05, **p<0.01, ***p<0.001; paired-sample t-test (against JE2).

In eukaryotic systems, glutathione acts as an important cellular antioxidant and contributes to varied physiological processes including metal homeostasis and alleviation of lipid oxidation stress, ultimately contributing to immune cell function (Dasgupta et al., 2010; Lushchak, 2012). We, therefore, wondered whether bacterial sequestration of glutathione may serve as a strategy to disrupt healthy phagocyte function and thereby promote bacterial survival. On average, we found no differences in cytotoxicity between the tested strains at 4 hours post-infection (Figure S5A). In contrast, we observed that THP1 cells infected with the GSH transporter-null double-mutant strain demonstrated increased cell death at 8 hours relative to the wild-type (Figure S5B). To investigate this further, we quantified the production of inflammatory cytokines at 6 hours post-infection, representing the mid-point between these two states. At this time point, cells infected with the double-mutant strain produced significantly lower levels of both IL-1β and IL6, though production of TNFα was seemingly unaffected (Figure S5C). While preliminary, these observations suggest that glutathione transporters allow *S. aureus* to modulate the macrophage pro-inflammatory status and maintain the intracellular niche by ultimately prolonging host-cell survival.

### Glutathione transport is not required during systemic murine infection

Our data indicates a role for bacterial GSH uptake in intracellular environments, but the contribution of this process toward fitness during infection is not yet understood. Previous work has demonstrated that disruption of *gis* does not significantly attenuate *S. aureus* JE2 during systemic infection in a murine model (Lensmire et al., 2023). Such an observation may be explained by the capacity of DtpT to facilitate GSH utilisation at concentrations encountered *in vivo*. To determine whether *dtpT* contributes toward fitness in this model, we carried out systemic infections with wild type JE2 and an isogenic mutant of *dtpT*, as well as a *dtpT*/*gis* double mutant strain (see supplemental methods SM1). In single-strain infection experiments, neither mutant demonstrated a pronounced difference in bacterial burden in the heart or kidneys at 96 hours post-infection. We did, however, observe significant attenuation of the *dtpT* mutant strain in the liver compared to the wild type at this time point (Figure S6A), though we failed to observe this phenotype for the *dtpT*/*gis* double mutant strain (Figure S6B). In a competitive infection assay, wild type JE2 did not outcompete the *dtpT* mutant strain in the liver (competitive index ≈ 1) but did outcompete a *gisB* mutant (Figure S6C). The competitive index of JE2 was greater against the *dtpT*/*gis* double-mutant strain on-average when compared to either of the *dtpT* or *gisB* single mutants, though this trend was not statistically significant (Figure S6C). Ultimately, our data suggest that glutathione uptake may contribute toward bacterial survival in the liver, though alternative nutrient sulphur sources available to the bacterium are seemingly able to compensate for the inability to utilise GSH in this environment.

### Recognition of GSH by DtpT in a vertical conformation during transport

Based on our discovery of physiologically relevant GSH transport through DtpT, we next sought to understand more about how this important tripeptide is recognised during transport. Previously, we demonstrated how the peptide-conjugated thioalcohol S-[1-(2-hydroxyethyl)-1-methylbutyl]-L-cysteinylglycine (S-Cys-Gly-3M3SH) is accommodated in an unconventional vertical conformation during transport by the *Staphylococcus hominis* POT PepT_Sh_ (Minhas et al., 2018). Given the close sequence homology between this protein and DtpT (87% sequence identity; including all residues previously implicated in substrate binding) (Figure S7) and the conserved L-Cys-Gly dipeptide backbone common to both substrates, we hypothesised that GSH and S-Cys-Gly-3M3SH may share a common binding mode in DtpT/PepT_Sh_ during transport.

To investigate this further, we generated a structural model of DtpT in the inward-open conformation based on the existing PepT_Sh_ complex structure (accession: 6EXS) and the binding mode of GSH in this protein was predicted using the protein-ligand docking tool CB-DOCK 2 (Figure 7A-D). In the predicted conformation, the L-Cys-Gly group of GSH occupies a similar position to the equivalent group in the S-Cys-Gly-3M3SH structure, albeit shifted slightly closer to the N-terminal bundle of the protein such that the γ-glutamyl group is fully accommodated within the extended 3M3SH-binding pocket (Figure 7D). In this position, the glutamate amino group is coordinated through polar contacts with the sidechain of Gln344 and the backbone carbonyl of Phe343, while the sidechain of Gln310 sits within hydrogen bonding distance of the glutamate α-carboxyl group (Figure 7C). Glu311 also sits adjacent to the substrate thiol group and may be involved in stabilising this group via hydrogen bonding or electrostatic interactions. For both substrates, the glycyl carboxyl terminus is anchored via a hydrogen bond with the sidechain of Tyr41 and sits adjacent to Asn167, both of which are conserved across diverse POT proteins (Figure S7). It is worth noting that the docked GSH conformation shown here does not take into account the presence of water molecules in the binding cleft, which were previously implicated in coordinating the L-Cys-Gly moiety of S-Cys-Gly-3M3SH (Figure 7B) and may similarly interact with the substrate in the proposed GSH binding conformation (as depicted in Figure 7C).

**Figure 7.**
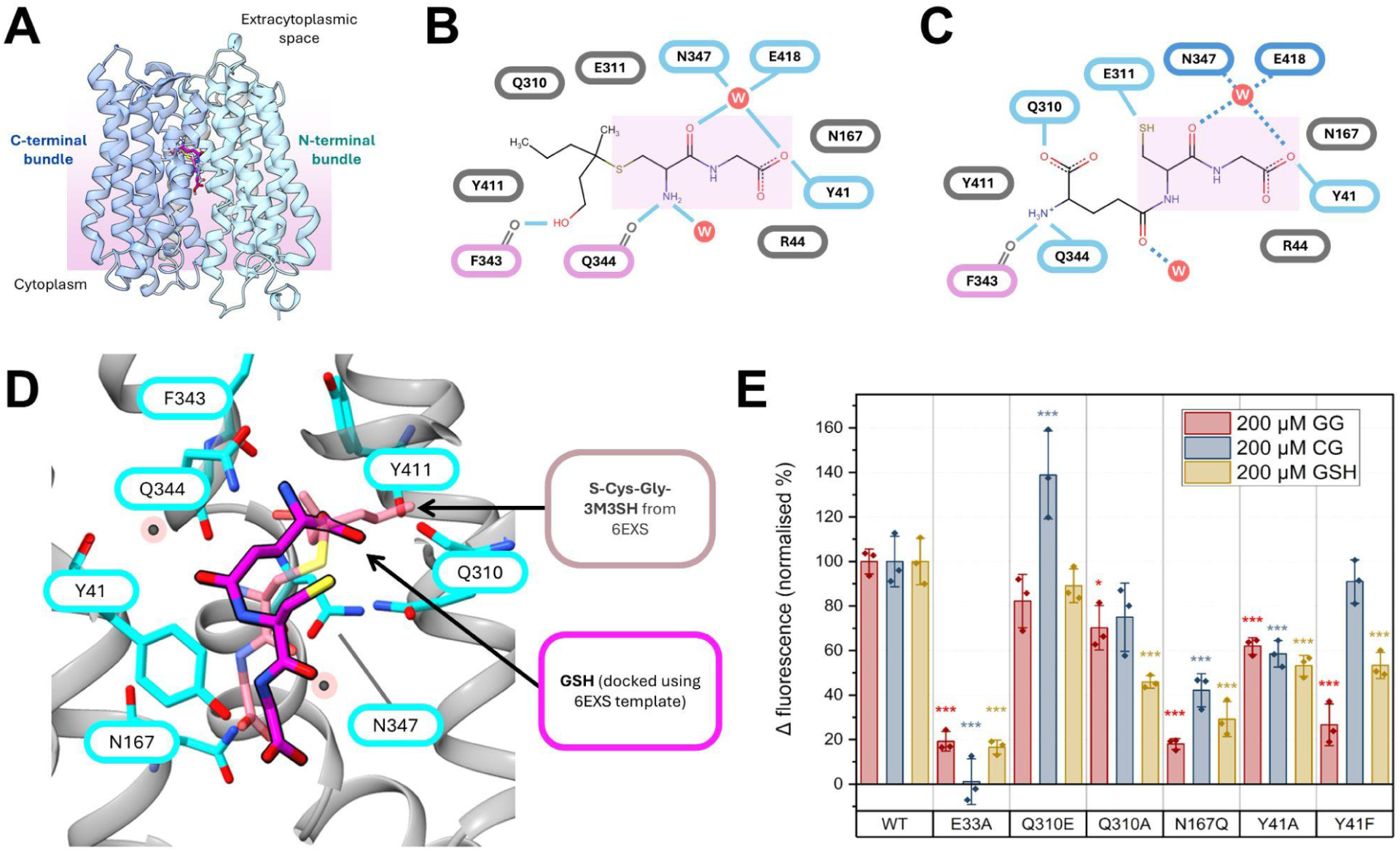
Molecular docking predicts a vertical binding mode for GSH in DtpT. (**A**) A structural model of DtpT in the inward-open conformation was generated with high confidence (QMEANDisCo Global = 0.87 ± 0.05, suggesting high per-residue quality for the model against the PepT_Sh_ template). GSH (pink) was docked into the DtpT model using the existing S-Cys-Gly-3M3SH binding conformation (light pink) as a template (docking score = -5.0 kcal/mol). (**B-C**) Diagrammatic overview of protein-ligand interactions underpinning substrate coordination between PepT_Sh_ and S-Cys-Gly-3M3SH (**B**) and the equivalent interactions between DtpT and GSH (**C**) in the proposed conformation. Solid blue lines indicate hydrogen bonds and polar contacts with sidechains (blue border) or main-chain groups (pink border). Dashed, dark-blue lines in (**C**) indicate possible contacts with water molecules (W). (**D**) Zoomed-in view of the proposed GSH binding mode in DtpT (grey ribbons). The known S-Cys-Gly-3M3SH binding conformation is shown for comparison. Residues surrounding the predicted binding site are shown as sticks (cyan) and labelled. (**E**) Transport data for DtpT mutant variants against GSH and two dipeptides, expressed as a percentage of the activity for the wild type (WT) protein. Bars indicate the mean ± standard deviation. *p < 0.05, ***p < 0.005; one-way ANOVA with Tukey honestly significant difference (HSD) test (against WT).

In order to assess the validity of this predicted binding model, mutant variants were generated with alterations in residues predicted to be involved in GSH binding and transport (Figure 7E; Figure S8). Glu33 forms part of the conserved E^33^xxERFxYY^41^ motif found ubiquitously in POT transporters and is known to contribute toward proton coupling during the transport cycle (Aduri et al., 2015; Lichtinger et al., 2024; Solcan et al., 2012). Unsurprisingly, an E33A mutant was found to be completely inactive against GSH and two dipeptide substrates, consistent with previous results in other POT family members (Figure 7E; Figure S8). Substitution of Tyr41 to either alanine or phenylalanine caused a significant decrease in GSH transport, consistent with the expected role of this residue in coordinating the substrate carboxy terminus. Similarly, substitution of Asn167 to glutamine strongly attenuated transport for all tested substrates, presumably by occluding the binding site (Figure 7E). GSH transport was disproportionately attenuated in a Q310A mutant variant when compared to either Gly-Gly or Cys-Gly, supporting the proposed vertical binding conformation of this tripeptide (Figure 7E). Conversely, substitution of this residue for glutamate (Q310E) did not significantly attenuate GSH transport (Figure 7E). Overall, our findings suggest that that GSH may be accommodated by DtpT in a similar conformation to S-Cys-Gly-3M3SH in the existing PepT_Sh_ complex structure, and this may indicate a unique mechanism by which POTs recognise sulphur-containing tripeptides.

### POT-mediated glutathione transport may be conserved in other Gram-positive intracellular pathogens

To the best of our knowledge, our findings represent the first evidence of POT-mediated GSH transport in a bacterium. Whether or not this function is conserved in the wider POT superfamily remains unclear, especially given that Gln310 (or Glu in the equivalent position) is seemingly required for strong transport of GSH (Figure 7E), but this feature is not well conserved across alternative POT proteins studied to-date (Figure S7).

One exciting possibility is that POT-mediated GSH transport is a conserved trait utilised by Gram-positive bacteria to assimilate nutrient sulphur in intracellular environments. *L. monocytogenes* is perhaps the most well characterised Gram-positive intracellular bacterial pathogen, and glutathione acquisition is a known determinant of fitness in this bacterium (Berude et al., 2024; Reniere et al., 2015). Similarly to *S. aureus*, *L. monocytogenes* possesses an ABC glutathione import system, but mutants of this system maintain the ability to utilise exogenous GSH at millimolar concentrations (Berude et al., 2024). *L. monocytogenes* also possesses a poorly characterised POT protein (LMON_0555 in *L. monocytogenes* EGD; Figure 8A) with approximately 50% identity to DtpT, and this protein has previously been associated with fitness in a murine liver infection model (Wouters et al., 2005). Notably, all residues predicted to contribute toward GSH binding in DtpT are conserved in LMON_0555, with the exception of Gln310 which is substituted for a Glu residue in the equivalent position (Figure 8B). In DtpT, this substitution did not significantly affect GSH transport (Figure 7E). Hence, we speculate that LMON_0555 is likely to serve as a secondary glutathione acquisition system in *L. monocytogenes*, and this activity may be relevant during bacterial survival *in vivo*.

**Figure 8.**
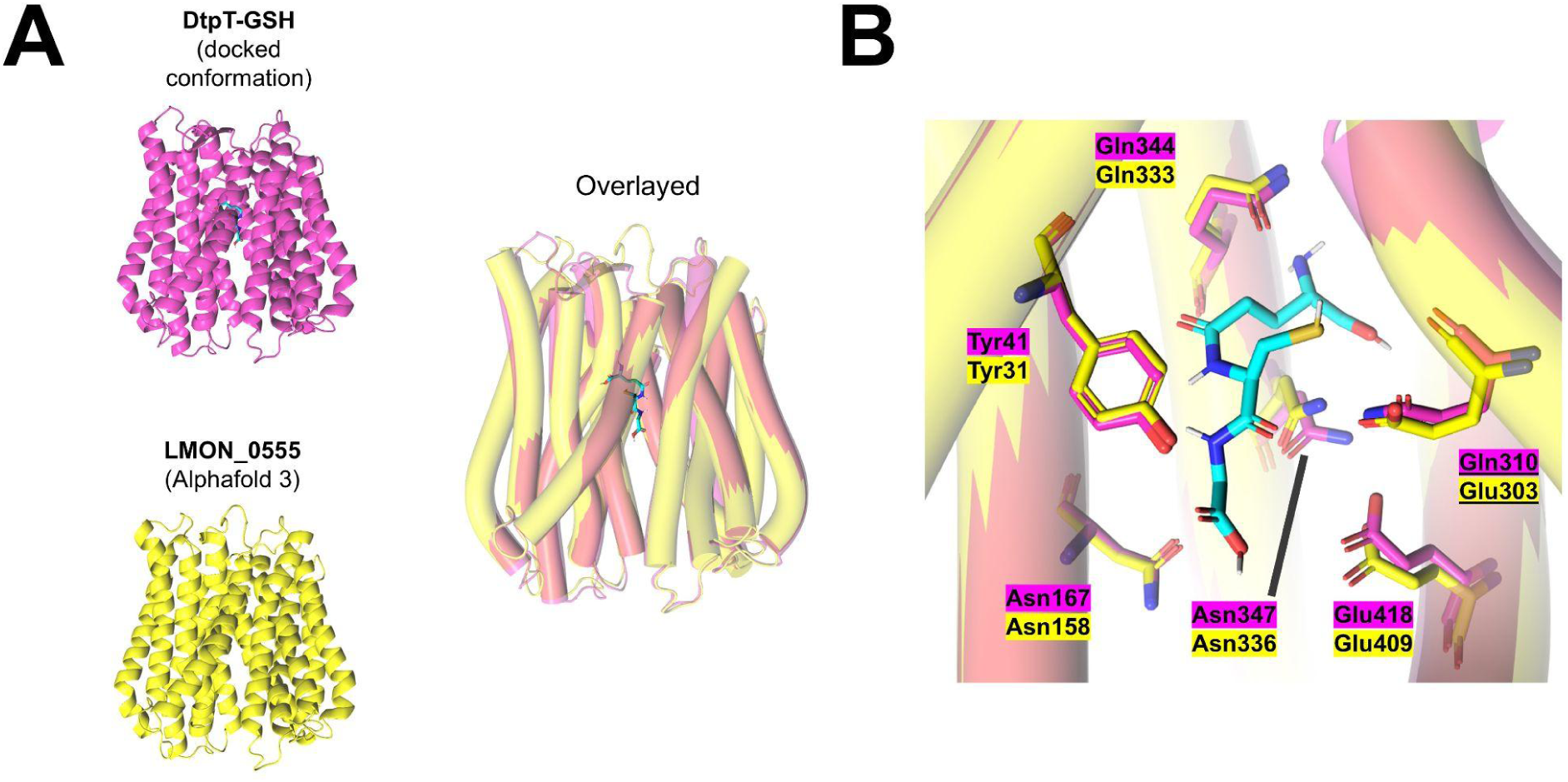
AlphaFold 3 predicts conservation of the proposed GSH binding site between DtpT and LMON_0555. (**A**) (left) Comparison of the previously generated structural model of DtpT (pink) with an AlphaFold 3 predicted structure of LMON_0555 (Yellow). The predicted binding conformation of GSH (cyan) in DtpT is also shown. (right) Overlaying these structures reveals a high degree of structural conservation (helices are depicted as cylinders to facilitate structural comparison). (**B**) Residues forming the proposed GSH binding site in DtpT and equivalent residues in LMON_0555 are shown as sticks and labelled in each case.

## Discussion

The promiscuous nature of peptide transport proteins and their functional redundancy in many organisms have made these systems historically challenging to characterise regarding substrate range and preference. Molecular dynamics simulations and similar computational methods have been combined with biochemical and structural methods in order predict substrate preferences in POTs, but experimental evidence for the applicability of these models to biological systems is lacking (Kotov et al., 2023; Li et al., 2022; Samsudin et al., 2016). Here, we present an extensive cell-based analysis of peptide transporter function in *S. aureus* and apply biochemical analysis of DtpT to both verify and expand our functional understanding of this transporter. Our findings demonstrate that DtpT is highly promiscuous, evidenced by the considerable heterogeneity observed among putative transport targets identified here. While earlier work has focussed on the contribution of Opp3 toward the utilisation of host-derived peptides (Lehman et al., 2019), our data instead highlights how DtpT serves as a major route of uptake for diverse dipeptides and tripeptides in *S. aureus*, including GSH. Indeed, this broad substrate range may underpin the essentiality of DtpT previously observed in various animal infection models (Coulter et al., 1998).

A number of recent studies have demonstrated how exogenous amino acids play diverse functional roles during *S. aureus* infections, serving both as key nutrient sources and environmental signals with defined regulatory consequences (Shibamura-Fujiogi et al., 2022; Freiberg et al., 2024; Urso et al., 2024). However, such studies generally do not consider the potential contribution of host-derived oligopeptides as reservoirs for these amino acids in biological environments. Our observations reveal a novel degree of substrate selectivity within the peptide transport systems on *S. aureus*, which may point toward defined roles for these systems in accumulation of specific amino acids during growth.

For instance, glutamate is known to serve as a major gluconeogenic carbon source for *S. aureus* and strains lacking the glutamate dehydrogenase GudB are attenuated under glucose-limited conditions (Halsey et al., 2017). Glutamate restriction has pleiotropic effects on cellular physiology, ultimately resulting in increased biofilm formation *in vitro* and *in vivo* (Shibamura-Fujiogi et al., 2022). Taken together, these observations implicate exogenous glutamate as an important factor contributing to both growth and environmental sensing in *S. aureus*. Acquisition of exogenous glutamate in *S. aureus* is primarily facilitated by the amino acid permease GltS (Zeden et al., 2020). Here, we demonstrate that DtpT is responsible for the utilisation of diverse glutamate-containing dipeptides and show that these peptides are specifically enriched among DtpT substrates (Figure 3A). These findings suggest that DtpT-mediated peptide uptake may serve as an additional source of exogenous glutamate in peptide-rich growth environments.

Our PM data reinforces the fact that DtpT serves as the primary route of dipeptide uptake in *S. aureus*, but also reveals some overlap in substrate specificity between this transporter and Opp3. The identification of a subset of peptides with redundant routes of internalisation serves as evidence that these peptides are of high value to the cell. Specifically, our data demonstrates how diverse arginine-containing dipeptides are utilised by *S. aureus* via either DtpT or Opp3 (Figure 2; 3B). *S. aureus* USA300 rapidly internalises arginine during growth but converts neither proline nor glutamate to arginine when grown *in vitro*, making exogenous arginine essential for growth (Halsey et al., 2017; Reslane et al., 2022). Experimental evidence has demonstrated how arginine restriction stimulates antibiotic tolerance in *S. aureus* biofilms, implicating arginine availability as an important environmental stimulus modulating *S. aureus* cell behaviour (Freiberg et al., 2024). We propose that the redundancy observed here for uptake of Arg-containing peptides likely signifies an evolutionary strategy to increase the diversity of potential arginine sources available to the cell, which agrees with the apparent importance of exogenous arginine in *S. aureus* physiology. Furthermore, the presence of two Arg-peptide transporters with varying substrate affinities in *S. aureus* may facilitate the utilisation of these peptides in growth environments with variable Arg-containing peptide availability.

In a similar vein, DtpT is capable of transporting the glutamine-containing dipeptide Ala-Gln (Figure 4C), but our PM data points toward the existence of a yet-unidentified additional utilisation system which is specific for glutamine-containing peptides (Figure 3A). The Opp4 transport system is an uncharacterised Opp-homologue which is widespread in *S. aureus* and is likely to have been generated via duplication of Opp3 (Yu et al., 2014). Given their close evolutionary relationship, we propose that Opp4 is likely to be a peptide transporter like Opp3 and is, therefore, a strong candidate to fulfil the role of the alternative Gln-peptide transport system in *S. aureus*. While ACME Opp may also contribute toward peptide transport, this system is more closely related to Opp1/2 and is, therefore, more likely to play a role in metal ion transport. Overall, the existence of an alternate utilisation pathway for Gln-containing peptides may point towards a yet underappreciated role for exogenous glutamine in *S. aureus* physiology.

In contrast to the examples listed above, we demonstrate that tyrosine-containing dipeptides are scarcely utilised by *S. aureus* (Figure 3C). This is perhaps unsurprising, given that *S. aureus* JE2 lacks a complete catabolic pathway for tyrosine utilisation and does not rapidly consume exogenous tyrosine when grown in defined media (Halsey et al., 2017). Intriguingly, no DtpT-mediated transport was observed for the sole tyrosine-containing peptide (Val-Tyr) assessed here (Figure 4B-C). Our findings therefore suggest that DtpT possesses a distinct biochemical preference toward peptide containing readily catabolised amino acids over those which lack the potential to serve as a nitrogen source to fuel growth. That being said, we do observe the utilisation of some dipeptides exclusively composed of the branched-chain amino acids (i.e. leucine, isoleucine and valine) and demonstrate that several such peptides are utilised via DtpT, despite the lack of a known catabolic pathway for these amino acids in JE2 (Halsey et al., 2017). For instance, we demonstrate that Leu-Leu is utilised robustly by JE2 and belongs to cluster 2, implicating both DtpT and Opp3 in the transport of this peptide (Figure 2). Our findings, therefore, suggest that branched chain amino acids derived from short peptides can serve as a nitrogen source for JE2, though the catabolic pathways involved remain to be elucidated.

Recent evidence points toward an essential role for amino acids (specifically, collagen-derived proline) in skin colonisation (Lehman et al., 2023; Urso et al., 2024), and it has previously been suggested that host-derived peptides serve as an important source for these amino acids during infection (Lehman et al., 2019). While we do not observe a clear preference for Pro-containing peptides among DtpT or Opp3 substrates, we do demonstrate preferable transport of Gly-Pro by DtpT in our transport assay (Figure 4C). Hence, our findings implicate DtpT in the utilisation of a collagen degradation product, and demonstrate how DtpT may contribute toward proline accumulation *in vivo*.

Glutathione is a ubiquitous metabolite in eukaryotic systems and is the predominant thiol present in intracellular niches, as well as being enriched in other environments associated with *S. aureus* infection including the sputum of cystic fibrosis patients (Shull et al., 2024). In most biological contexts, GSH is the most abundant form of glutathione by a significant margin (Hansen et al., 2009). Here, we identify GSH as a substrate of DtpT and demonstrate that either DtpT or Gis-mediated transport is sufficient to meet the nutrient sulphur requirements of *S. aureus* at physiologically relevant GSH concentrations (Figure 5). Generally, binding-protein dependant transport systems (including ABC transporters) operate with high affinities, whilst POTs and similar MFS systems have been reported as having substrate affinities in the high-micromolar or low millimolar range (Kotov et al., 2023; Samsudin et al., 2016). We, therefore, predict that Gis may serve as the dominant system involved in GSH scavenging under conditions where the metabolite is scarce, whilst both systems likely contribute to transport in GSH-enriched environments. Indeed, the phenotypes associated with our mutant strains support such a model (Figure 5C; Figure S4).

Finally, we identify *S. aureus* glutathione transporters as important fitness determinants during intracellular survival inside macrophage cells (Figure 6), representing an environment in which GSH is present at millimolar concentrations (Meister, 1988). In *Mycobacterium tuberculosis* (Mtb), glutathione transport has been implicated in regulating the activity of Mtb-infected macrophages and thereby contributing toward intracellular survival and immune evasion (Dasgupta et al., 2010). Our macrophage infection data similarly demonstrates how the presence of glutathione transporters affects both bacterial survival (Figure 6) and host-cell behaviour, seemingly allowing the pathogen to modulate innate immune signalling responses and subvert premature host-cell death (Figure S5). In *L. monocytogenes*, the major glutathione transporter (Ctp) has similarly been linked to virulence in a murine infection model (Berude et al., 2024), and our structural modelling provides compelling evidence that GSH transport is also a conserved function of the *L. monocytogenes* POT transporter (Figure 8). Hence, GSH transport may represent a conserved strategy utilised by bacterial pathogens in order to enable intracellular survival, thereby avoiding exposure to extracellular stresses and immune clearance. More work is now needed to better understand how host-pathogen interactions are affected as a result of bacterial glutathione transport and determine whether these responses vary in different bacteria and host-cell types.

Overall, this work has significantly expanded our knowledge of peptide utilisation in *S. aureus* and shed light on a novel role for glutathione transport in host-pathogen interactions during intracellular infection. However, with the diversity of roles ascribed to peptide transporters in other organisms – including cell-wall turnover (Maqbool et al., 2011; Park et al., 1998; Simpson et al., 2023), antimicrobial peptide resistance (Parra-Lopez et al., 1993; Rivera et al., 2024) and cell signalling (Dubois et al., 2019; Nasher et al., 2018; Zheng et al., 2018) – it is clear that this is an area of *S. aureus* biology where much remains to be discovered and understood.

## Methods

### Bacterial strains and growth conditions

All *S. aureus* strains used in this study were routinely cultured in tryptic soy broth (TSB) with shaking or on solid tryptic soy agar (TSA) medium at 37°C. *E. coli* strains were routinely cultured in lysogeny broth (LB) with shaking or on LB agar at 37°C. Selective antibiotics were added during growth when appropriate at the following concentrations: 50 µg/ml kanamycin, 100 µg/ml ampicillin, 10 µg/ml erythromycin, 10 µg/ml chloramphenicol.

*S. aureus* JE2 transposon mutants were acquired from the Nebraska Transposon Mutant Library (NTML) (Fey et al., 2013) and transduced into wild-type JE2 via Ф85 bacteriophage in order minimise off-target genetic inconsistencies between strains. To generate double mutant strains, the Erm^R^ resistance cassette was exchanged with an unmarked sequence using the allelic exchange vector pTnT, as described previously (Bose et al., 2013). Plasmids were generated using the ClonExpress II One Step Cloning Kit (Vazyme) according to manufacturer’s instructions. DtpT complementation in *S. aureus* mutant strains was achieved via introduction of the entire *dtpT* locus into the *E. coli* / *S. aureus* shuttle vector pSK5630 modified by the addition of the native *dtpT* promoter sequence (herein referred to as pSK56_pT_) (Grkovic et al., 2003).

A full list of bacterial strains utilised in this work are provided in supplemental table ST1. Primers utilised for diagnostic PCR, cloning and sequencing of genetic elements are provided in supplemental table ST2.

### Phenotype microarrays and analysis

Peptide utilisation by *S. aureus* strain JE2 and mutant derivatives was assessed on phenotype microarray plates PM 6-8 (Biolog) using a modified version of the manufacturer’s protocol. Cells grown on TSA overnight were transferred from solid media into 3 ml IF-0α buffer using a sterile cotton swab and pelleted. Cells were washed twice and resuspended in IF-0α. This cell suspension was then diluted to a final OD_600_ of 0.004 in 9.975 ml IF-0α and supplemented with 110 µl redox dye H (Biolog) and 915 µl 12X PM additive solution (240 mM tricarballylic acid, pH 7.1; 24 mM MgCl_2_; 12 mM CaCl_2_; 150 µM L-cystine, pH 8.5; 0.06% w/v yeast extract; 0.06% v/v tween 80; 6 mM D-glucose; 12 mM pyruvate). PM plates were inoculated with 100 µl/well of the prepared cell suspension and incubated for 36 hours at 37°C with continuous slow shaking in an Epoch2 microplate spectrophotometer (Agilent). OD_590_ and OD_750_ were measured every 15 minutes.

Signal curves (OD_590_ - OD_750_ against time) for all four strains were grouped for each peptide and then analysed by Uniform Manifold Approximation and Projection (UMAP) using the UMAP R library and a random seed of 888 (McInnes et al., 2018). K-means clustering was used on the same grouped data with "kmeans" R function and k=7. Empirical area under the curve (eAUC) was calculated using Growthcurver (Sprouffske & Wagner, 2016).

### Peptide utilisation assays

Utilisation of specific peptides was assessed using glucose-supplemented chemically defined medium (CDMG) based on a previously reported methodology (Hussain et al., 1991). The complete composition of the CDMG used here is provided in supplemental table ST3. For assessing the utilisation of arginine-containing peptides, L-arginine was omitted from the medium (CDMG - *R*). For assessing the utilisation of glutathione, L-cysteine and L-methionine were omitted from the medium and (NH_4_)_2_SO_4_·FeSO_4_·6H_2_O was substituted for 15 µM iron (III) citrate (CDMG - *s*). Reduced glutathione solutions were prepared under anaerobic conditions to prevent changes in the oxidation state before commencing assays. Overnight cultures of *S. aureus* strain JE2 and mutant derivatives were diluted 1:100 into fresh TSB and grown to mid-exponential phase. Cells were then pelleted and washed twice in the assay medium (CDMG - *R* or CDMG - *s*) before being resuspended and diluted to a final OD_600_ of 0.05 in peptide-supplemented medium. 200 µl of each cell suspension was transferred to a Costar 96 well plate (Corning) and incubated for 24 hours at 37°C with shaking. OD_600_ was measured every 30 minutes.

For anaerobic growth assays, cultures were prepared as before being transferred to a Whitley A85 anaerobic workstation (Don Whitley Scientific) and diluted into assay medium which had been equilibrated under anaerobic conditions for at least 3 hours and supplemented with 100 mM sodium nitrate. Growth was monitored as above in a Stratus 96 well plate reader (Cerillo) for 18 hours at 37°C without shaking.

### Protein purification

The DtpT open reading frame was amplified directly from the *S. aureus* JE2 genome and cloned into the C-terminal octa-histidine/GFP-tagged fusion vector pWaldo (Drew et al., 2006) before being transformed into *E. coli* C43(DE3) (Miroux & Walker, 1996). Mutant variants were generated via site-directed PCR mutagenesis and blunt-end ligation with T4 DNA ligase (NEB) and all constructs were verified by sequencing.

Cells grown overnight in LB supplemented with kanamycin were diluted 1:100 into fresh media. Cultures were incubated at 37°C until an OD_600_ of 0.5 was reached. Expression was then induced by the addition of IPTG to a final concentration of 100 µM. Cultures were incubated overnight at 25°C with shaking. Cells were then pelleted, resuspended in 50 ml resuspension buffer (40 mM Tris, pH 7.5; 10% glycerol) and lysed by sonication. Crude lysates were clarified via centrifugation at 27,000 x g for 30 minutes. Membranes were then pelleted via ultracentrifugation at 164000 x g for 120 minutes. Membrane pellets were pooled and resuspended in 40 ml protein solubilisation buffer (40 mM Tris, pH 7.5; 200 mM NaCl; 20 mM imidazole; 10% glycerol; 0.5% n-dodecyl-B-pyromaltoside (DDM)) using a glass homogenizer before being incubated for 60 minutes at 4°C with rolling. The resultant suspension was then ultracentrifuged as before for 60 minutes to pellet leftover membranes and insoluble contaminants. Supernatant containing the solubilised membrane proteins was pooled and applied to a His-TRAP nickel affinity column (GE Healthcare) using an ÄKTA Protein Purification System (Cytiva). This column was washed with 50 ml column wash buffer (40 mM Tris, pH 7.5; 200 mM NaCl; 20 mM imidazole; 10% glycerol; 0.04% DDM) and the fusion protein was then eluted via application of column elution buffer (40 mM Tris, pH 7.5; 200 mM NaCl; 400 mM imidazole; 10% glycerol; 0.04% DDM).

Purified protein was exchanged into TEV reaction buffer (40 mM Tris, pH 7.5; 200 mM NaCl; 0.5 mM EDTA; 1 mM DTT; 10% glycerol; 0.04% DDM) using a HiTRAP desalting column (SLS). Protein was then mixed with TEV protease to a final ratio of approximately 3:1. The resultant reaction mixture was incubated overnight at 4°C. Pure DtpT was isolated by applying the reaction mixture to a His-TRAP nickel affinity column as before and washing with column wash buffer, this time collecting the flow-through fractions. Pure protein was stored at -70°C.

### Reconstitution into proteoliposomes and transport assays

Purified DtpT was exchanged into reconstitution buffer (20 mM Tris pH 7.5; 150 mM NaCl; 0.03% DDM) and diluted to a final concentration of 0.5 mg/ml. Liposomes (composed of POPE and POPG in a 3:1 ratio) were prepared and extruded 11 times each through pre-soaked 0.8 µm and 0.4 µm track-etched polycarbonate membranes (Whatman) before diluting to a final concentration of 10 mg/ml in liposome buffer (50 mM KPi, pH 7).

10 mg lipid mixture was gradually added to 200 µg protein at room temperature before being incubated on ice for 60 minutes. 100 µl SM2 Biobeads (Biorad) suspended in water were added to the lipid-protein mixture, which was then incubated for 1 hour at 4°C while rotating. A further 130 µl Biobeads were added and the mixture was incubated as before for a further 3 hours. Biobeads were pelleted and the lipid-protein mixture was transferred to a fresh tube. 160 µl Biobeads were added and the mixture was incubated as before for a further 16 hours. Biobeads were removed as before and proteoliposomes were pelleted via ultracentrifugation at 158 000 × g for 30 minutes. Proteoliposomes were then resuspended in liposome buffer to a final protein concentration of 0.5 mg/ml (assuming 100% reconstitution efficiency) and subjected to three freeze-thaw cycles before dialysing extensively against liposome buffer at 4°C. Proteoliposomes were pelleted as before and resuspended in liposome buffer before being stored at -70°C.

For transport assays, proteoliposomes were pelleted as before and resuspended in inside buffer (5 mM HEPES, pH 6.8; 120 mM KCl; 2 mM MgSO_4_) spiked with 1 mM pyranine. Proteoliposomes were subjected to seven freeze-thaw cycles before being extruded through a pre-soaked 0.4 µm track-etched polycarbonate membrane. Proteoliposomes were then pelleted at 20°C and resuspended in inside buffer. Excess pyranine was removed using a microspin G25 desalting column (Cytiva) according to the manufacturer’s protocol. Liposomes were finally pelleted again at 20°C and resuspended to a final protein concentration of 1 mg/ml. For each assay, 6 µg of proteoliposomes were diluted 1:100 into outside buffer (5 mM HEPES, pH 6.8; 120 mM NaCl; 2 mM MgSO_4_). Assays were performed in 1 ml clear plastic microcuvettes using a Fluoromax-4 spectrophotometer (Horiba) with a magnetic flee for stirring. Fluorescence of pyranine was measured via excitation at 460 nm and emission at 510 nm. Substrate was added to the reaction at t = 25 s and transport was induced by the addition of 1 µM valinomycin at t = 50 s. All curves are normalised before the addition of valinomycin (t = 45 s) to facilitate comparison.

### Macrophage culture and differentiation

THP1 cells (ATCC) were cultured in macrophages media (RPMI1640 supplemented with 10% Fetal Calf Serum and Glutamine). To obtain a macrophage-like differentiation state, THP1 were differentiated for 48 hrs in macrophage media supplemented with 50 ng/mL Phorbol 12-myristate 13-acetate (in DMSO, Sigma-Aldrich). PMA-containing media was replaced with fresh macrophage media and cells were rested overnight prior to infection.

Human primary macrophages were differentiated from human peripheral blood mononuclear cells (PBMCs) obtained from the NHS (University of York Biology ethic committee project DB202111) and differentiated in complete macrophage media supplemented with 50 ng/mL macrophage colony-stimulating factor (mCSF, Proteintech) over 7 days.

### Macrophage infection

Infection of macrophages and macrophage-like cell lines was carried out based on previously published methods (Alves et al., 2024). Overnight cultures of *S. aureus* strain JE2 and mutant derivatives were diluted 1:100 into fresh TSB and grown to mid-exponential phase. Cells were then pelleted and washed twice in ice-cold phosphate-buffered saline (PBS) before being resuspended to a final cell density of 5 x 10^7^ CFU/ml. For infection, 10 µl bacterial suspension was added to 50,000 hMDM or THP-1 cells (D.O.I. = 10) in 90 µl Opti-MEM reduced serum media (Thermo Fisher) and plates were centrifuged for 5 minutes at 500 x g to promote contact. After 30 minutes of incubation at 37°C, the medium was exchanged for Opti-MEM supplemented with 100 µg/ml gentamicin. The addition of gentamicin was designated t = 0 h. After a further 30 minutes of incubation this medium was removed and cells were then maintained in Opti-MEM supplemented with 5 µg/ml gentamicin (Gibc).

To compare bacterial internalisation and survival, cells at t = 0.5 / 8 h were incubated in deionised water for 10 minutes before being lysed by repeated pipetting. Suspensions were serially diluted in PBS, spotted onto TSA and incubated overnight at 37°C to determine viable CFU. Cytotoxicity was measured by quantifying the activity of lactate dehydrogenase (LDH) in the extracellular medium (Cytotox 96 non-radioactive cytotoxicity kit, Promega) and normalised against cells lysed by 0.04% Triton X-100. Cytokine levels in cell culture medium were determined at t = 6 h by utilising human IL 1-β, IL 6 and TNF-α ELISA assay kits (invitrogen) according to the manufacturer’s protocols.

### Structural modelling and docking

A structural model of DtpT was generated using SWISS-MODEL (Waterhouse et al., 2018) using the solved complex structure of the *S. hominis* PepT_Sh_ (6EXS) as a template. For predicting how DtpT recognises glutathione, blind docking was carried out using this projected structure in CB-Dock 2 (Liu et al., 2022). A high confidence predicted structure of LMON_0555 (pTM = 0.94) was generated using Alphafold 3 (Abramson et al., 2024). Projected and docked structures were visualised in ChimeraX (UCSF).

## Author contributions

**Imran Khan**: Conceptualization, Formal analysis, Investigation, Methodology, Data processing, Writing – original draft, review & editing. **Sandy MacDonald**: Formal analysis, Data processing. **Sigurbjörn Markusson**: Investigation**. Paige J. Kies** and **Cristina Kraemer-Zimpel**: Investigation, Animal experiments. **Callum Robson:** Investigation. **Joanne L Parker**: Conceptualization, Methodology. **Simon Newstead**: Conceptualization, Methodology. Writing – review & editing. **Dave Boucher**: Conceptualization, Investigation, Methodology, Writing – review & editing. **Neal D. Hammer**: Conceptualization, Investigation Methodology, Supervision, Animal experiments. **Marjan Van Der Woude**: Conceptualization, Methodology, Supervision, Project administration, Funding acquisition, Writing – review & editing. **Gavin H Thomas**: Conceptualization, Methodology, Supervision, Project administration, Funding acquisition, Writing – review & editing.

## Acknowledgements

We are extremely grateful to the Burgess family for the award of a gift to the University of York that supported a PhD studentship to IK. We acknowledge ongoing support of the GHT lab through BB/X003035/1. Work carried out in the NDH lab was supported by the National Institute of Allergy and Infectious Diseases of the National Institutes of Health, R01 AI139074. The defined transposon mutant library used in this study was provided by the Network on Antimicrobial Resistance in Staphylococcus aureus (NARSA) for distribution by BEI Resources, NIAID, NIH: Nebraska Transposon Mutant Library (NTML) Screening Array NR-48501. We thank Dr Rebecca M. Corrigan (ORCiD: 0000-0002-6031-1148) for supplying the wild-type JE2 and NTML strains utilised in this study, and for providing valuable technical guidance.

