## Supplemental figures and tables for "Defined roles for the *Staphylococcus aureus* POT transporter DtpT in di/tripeptide uptake and glutathione utilisation inside human macrophages"

**Supplemental figure 1. Arginine-containing dipeptides restore growth under arginine limitation only when a viable route of uptake is available.**

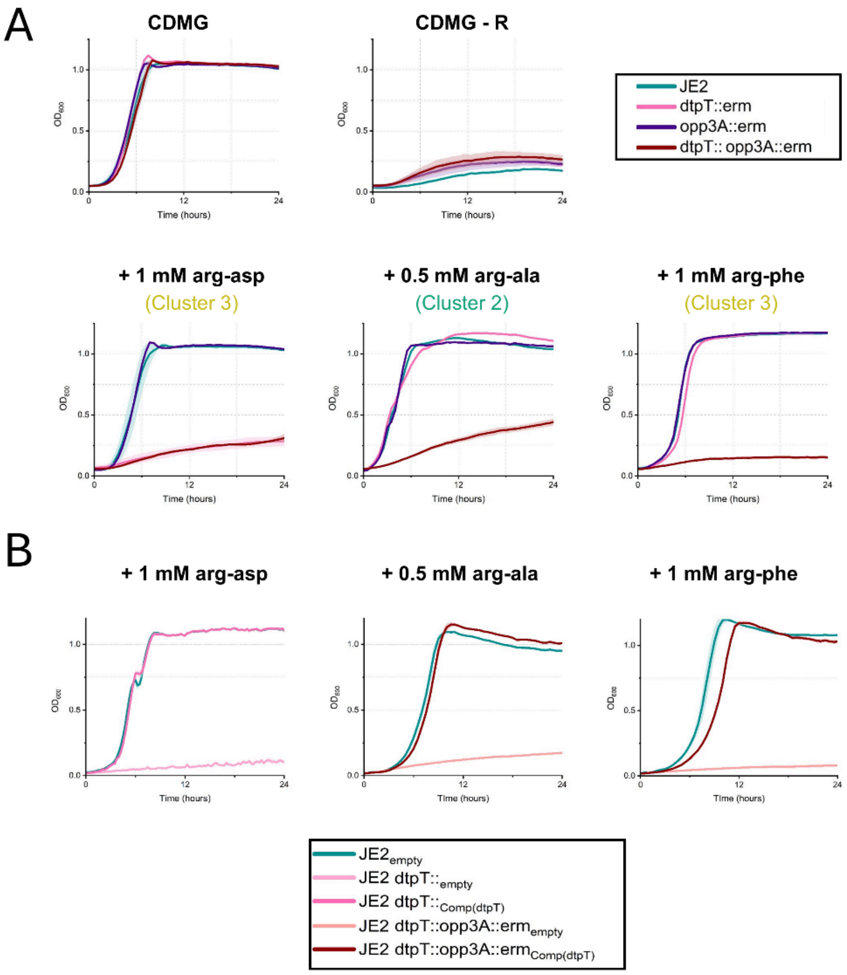

**Figure S1 – Arginine-containing dipeptides restore growth under arginine limitation only when a viable route of uptake is available.** In each case, strains were grown over 24 hours and $OD_{600}$ was measured every 30 minutes for three biological replicates. Curves indicate the mean values for each reading and the standard deviation in each case is indicated by the shaded area. (**A**) Growth of *S. aureus* strain JE2 and mutant derivatives in CDMG and CDMG – R. Growth of all strains is dependent on an exogenous source of arginine. (**B**) Growth of *S. aureus* strain JE2 and mutant derivatives in CDMG – R supplemented with three dipeptides, as labelled. Corresponding PM clusters are also provided. (**C**) Growth of deficient strains is restored by *in-trans* expression of *dtpT* under its native promoter in each case.

**Supplemental figure 2. Purification of DtpT from *E. coli* C43 (DE3)**

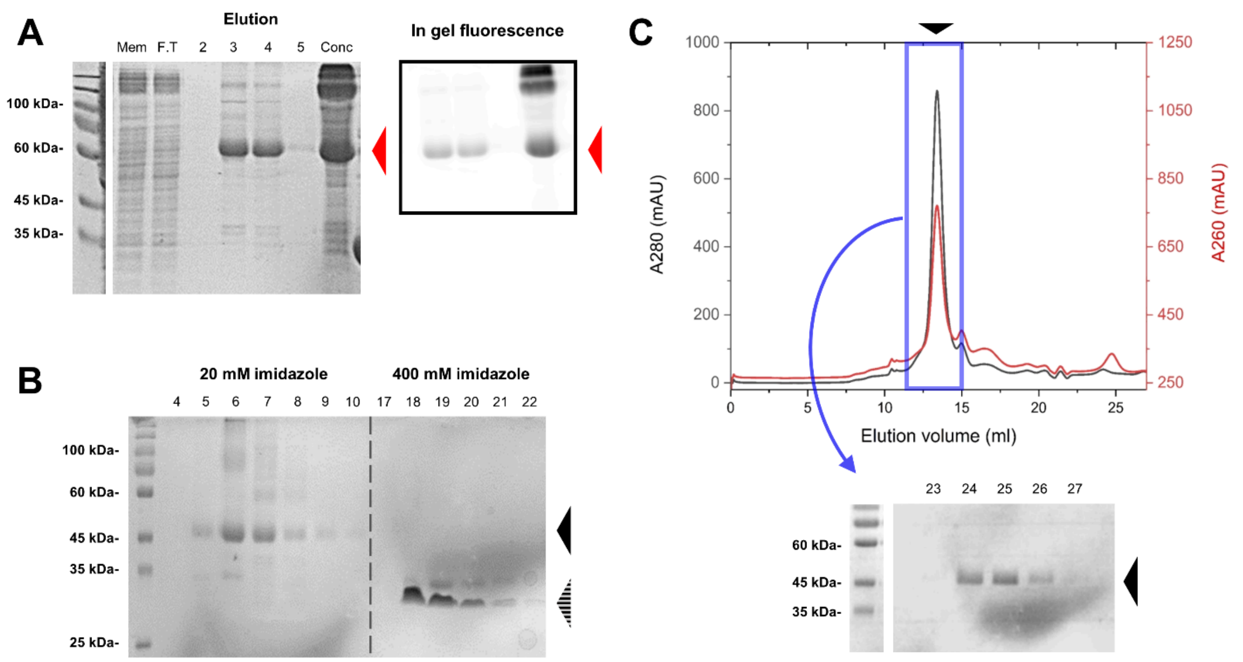

**Figure S2 – Purification of DtpT from *E. coli* C43 (DE3).** (**A**) Nickel affinity purification of a DtpT-GFP fusion protein. Images correspond to total protein (**left**; Coomassie blue stain) and DtpT-GFP only (**right**; in-gel GFP fluorescence). Mem = Membrane fraction. F.T = Column flow-through. Conc = concentrated fractions. (**B**) Representative gel image of a reverse nickel affinity purification of untagged DtpT after TEV cleavage of GFP-8His. (**C**) SEC purification of pure DtpT protein following TEV cleavage. A single sharp peak is seen at approximately 12.5 ml elution volume in the A280 trace (**upper**) corresponding to the pure DtpT protein, split across elution fractions 24 - 26 (**lower**). Overall, approximately 2.35 mg of pure DtpT protein was yielded from 2 L of bacterial culture as estimated from A280. The black arrow in each case indicates the position of a band corresponding to the pure DtpT protein. The red arrow indicates the position of a band corresponding to the expected DtpT-GFP fusion protein. The striped arrow indicates the position of sfGFP.

**Supplemental figure 3. DtpT proteoliposomes require both a valid substrate and potential gradient in order to drive proton-coupled transport.**

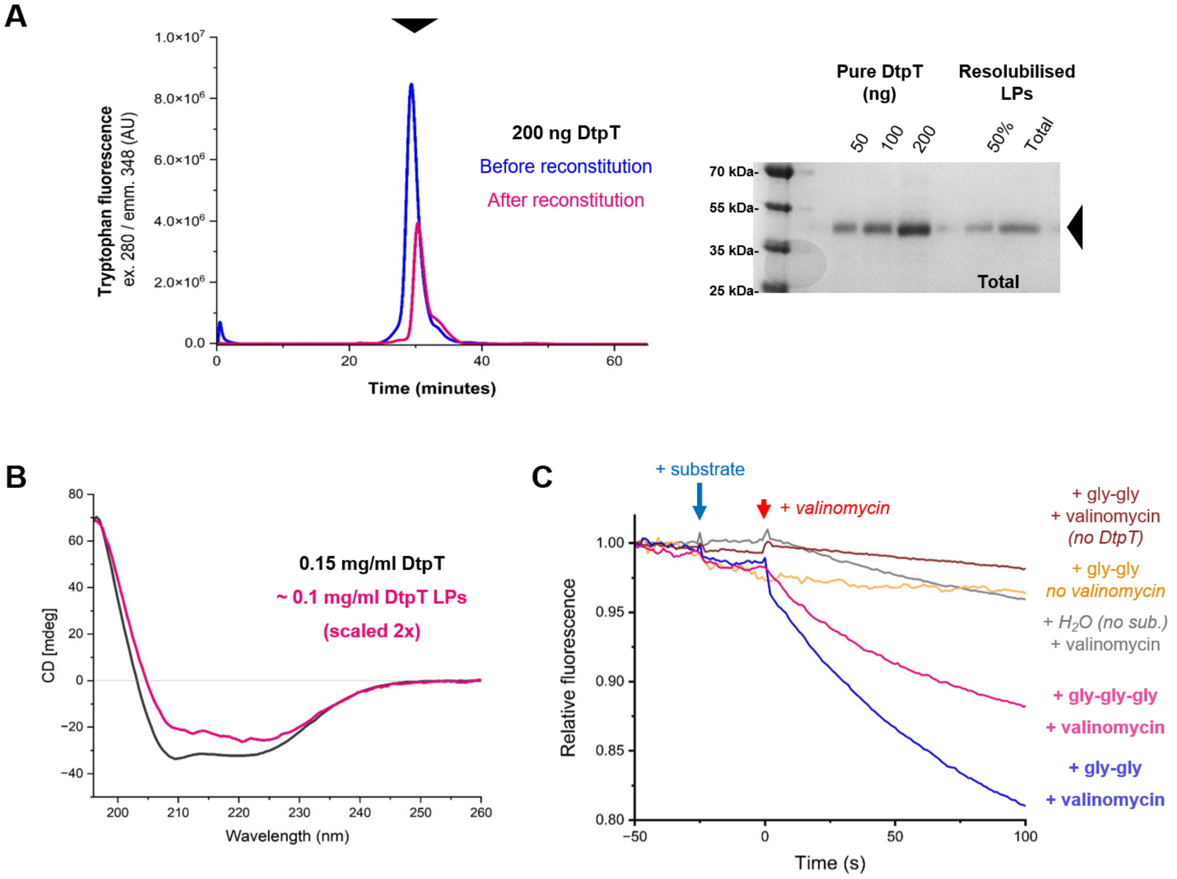

**Figure S3 – DtpT proteoliposomes require both a valid substrate and potential gradient in order to drive proton-coupled transport.** (**A**) Successful reconstitution of DtpT was confirmed by size exclusion chromatography (**left**) and SDS PAGE (**right**). The expected position of the soluble DtpT protein is indicated by the black arrow. (**B**) CD spectra of pure DtpT (grey) and DtpT liposomes (pink). A scaling factor of 2x has been applied to the DtpT-liposome spectrum to facilitate comparison with the detergent-soluble protein. (**C**) Validation of pyranine assays in DtpT liposomes. Curves show the mean of three replicates normalised to the initial fluorescence signal (t = -50 s). In each case, peptide (or $H_2O$) is added at t = -25 s (blue arrow) and a negative-inside $\Delta\Psi$ is established by the addition of valinomycin at t = 0 s (red arrow). No acidification is observed in the absence of DtpT. A slight decrease in fluorescence is observed upon addition of valinomycin in the absence of substrate, likely due to proton leakage. Strong acidification of the lumen indicative of DtpT-mediated transport requires both peptide substrate and $\Delta\Psi$.

**Supplemental figure 4. Extended growth assays provide further insight into the routes of glutathione transport in *S. aureus*.**

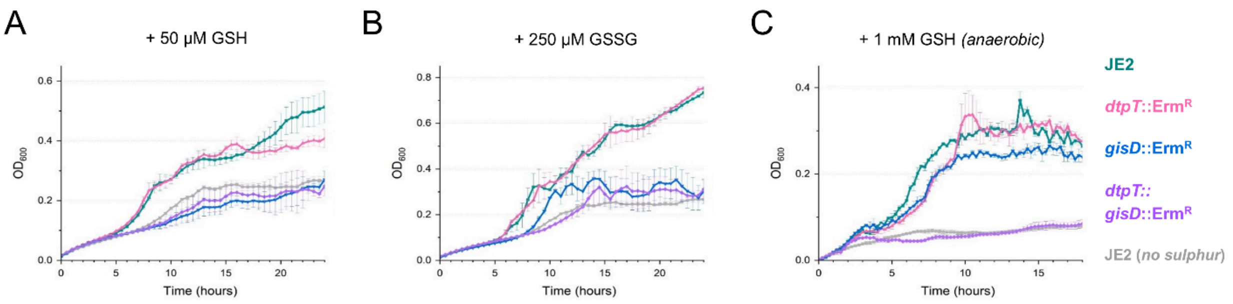

**Figure S4 – Extended growth assays provide further insight into the routes of glutathione transport in *S. aureus*.** Growth of *S. aureus* strain JE2 and mutant derivatives was assessed in CDMG – *s* supplemented with 50 µM GSH (**A**) or 250 µM GSSG (**B**) in aerobic conditions, as well as in the presence of 1 mM GSH under anaerobic conditions (**C**). For A and B, strains were grown over 24 hours and $OD_{600}$ was measured every 30 minutes. For C, strains were grown over 18 hours and $OD_{600}$ was measured every 15 minutes. Curves indicate the mean values for three biological replicates ± standard deviation.

**Supplemental figure 5. Glutathione transporters modulate the host response to infection inside THP-1 cells.**

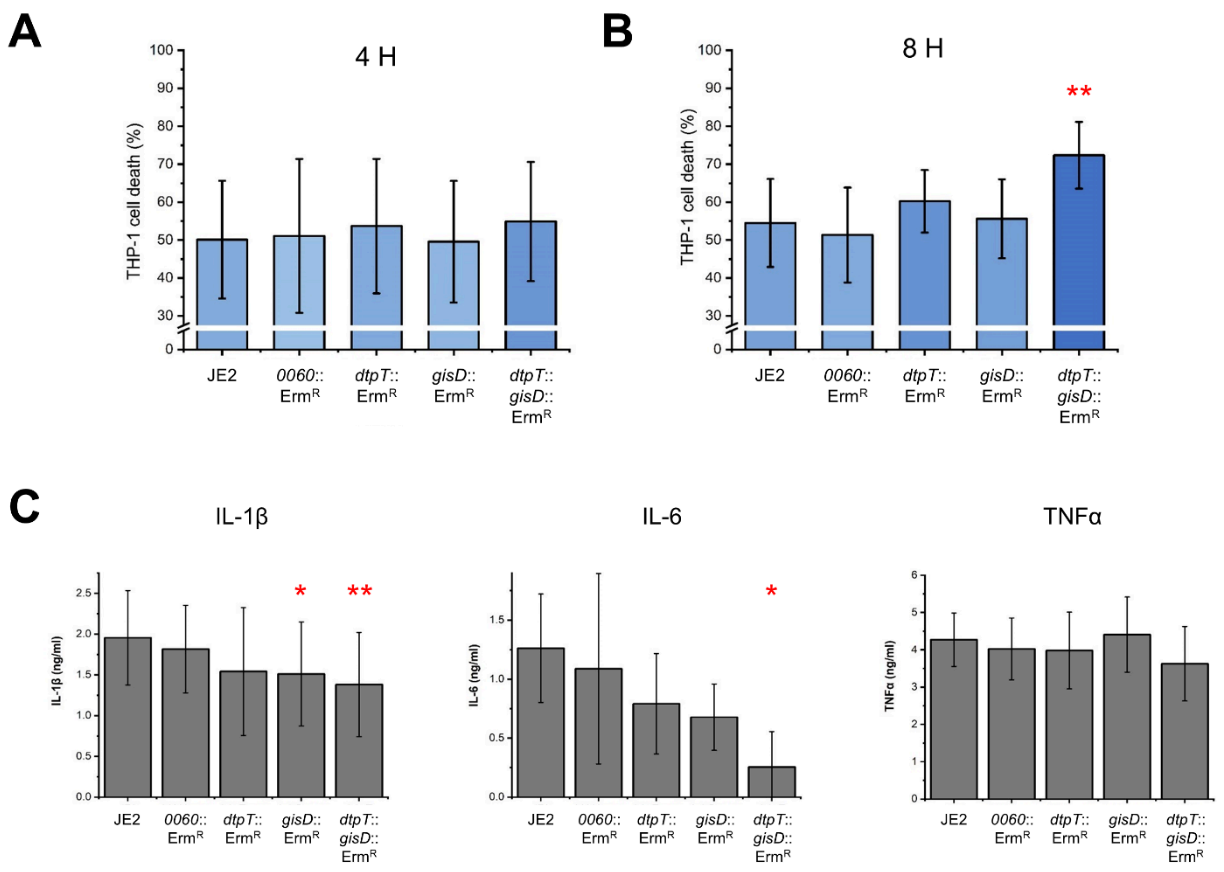

**Figure S5 – Glutathione transporters modulate the host response to infection inside THP-1 cells.** (**A - B**) Cell death as a result of bacterial infection was quantified at 4 hours (**A**) and 8 hours (**B**) post-infection for THP1 cells infected with JE2 or mutant derivatives (n = 6, each measured in technical triplicate). Cell death was quantified by measuring the activity of extracellular LDH in cell culture supernatant. Values are given as a percentage of the LDH activity for cells lysed by addition of 0.04% triton X-100 and corrected for background signal. (**C**) Production of inflammatory cytokines by infected THP-1 cells at 6 hours post-infection (n = 4, each measured in technical triplicate). *p<0.05, **p<0.01; as determined by paired-sample t-test (vs JE2).

**Supplemental figure 6. Virulence quantification of *dtpT*::Tn and Δ*gis dtpT*::Tn mutants in a murine model of systemic infection.**

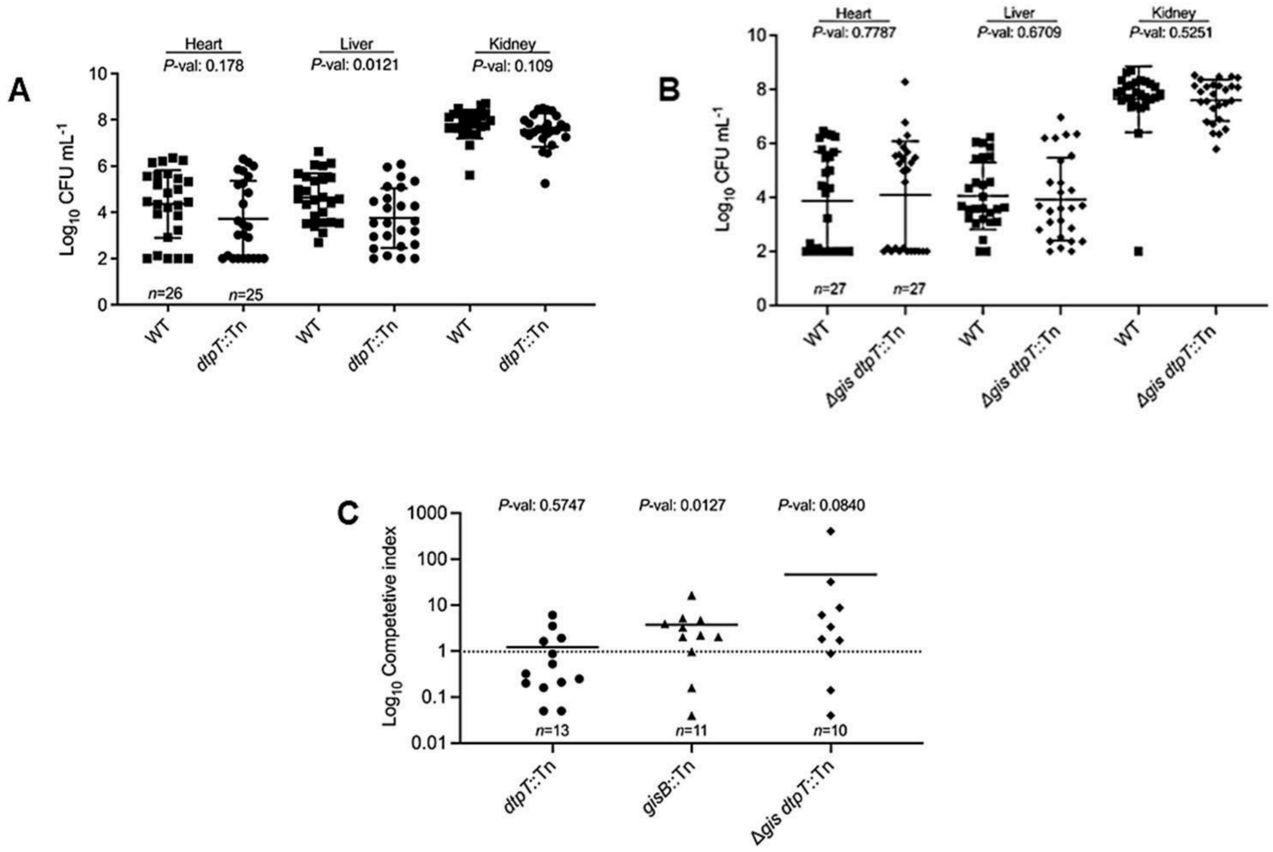

**Figure S6 – Virulence quantification of *dtpT*::Tn and Δ*gis dtpT*::Tn mutants in a murine model of systemic infection.** (**A - B**) Bacterial burdens within indicated organs after systemic inoculation of BALB/cJ mice were enumerated after 96 h of infection with either WT (squares), *dtpT*::Tn (circles) (**A**) or **Δ***gis dtpT*::Tn (diamonds) (**B**). Bacterial burdens are presented as $\log_{10}$ transformed CFUs $mL^{-1}$ for liver, combined kidneys, and heart. The mean and standard deviation are presented as horizontal lines. Normality was determined using a Shapiro-Wilk test. *p* values were determined by Mann-Whitney test. (**C**) Competitive indices (CI) for *dtpT*::Tn, *gisB*::Tn, and Δ*gis dtpT*::Tn were determined from the livers of Balb/cJ mice systemically inoculated with a 1:1 mixture of WT and the indicated mutant strain. The liver output ratio of WT to indicated mutant CFU $mL^{-1}$ in the liver over the input WT to mutant CFU $mL^{-1}$ were used to calculate the CI. The mean of WT:indicated mutant CI are presented as a horizontal bar and *p* values denoted for each competition were determined by Wilcoxon signed-ranked test.

**Supplemental figure 7. Multiple sequence alignment of DtpT and structurally characterised POTs**

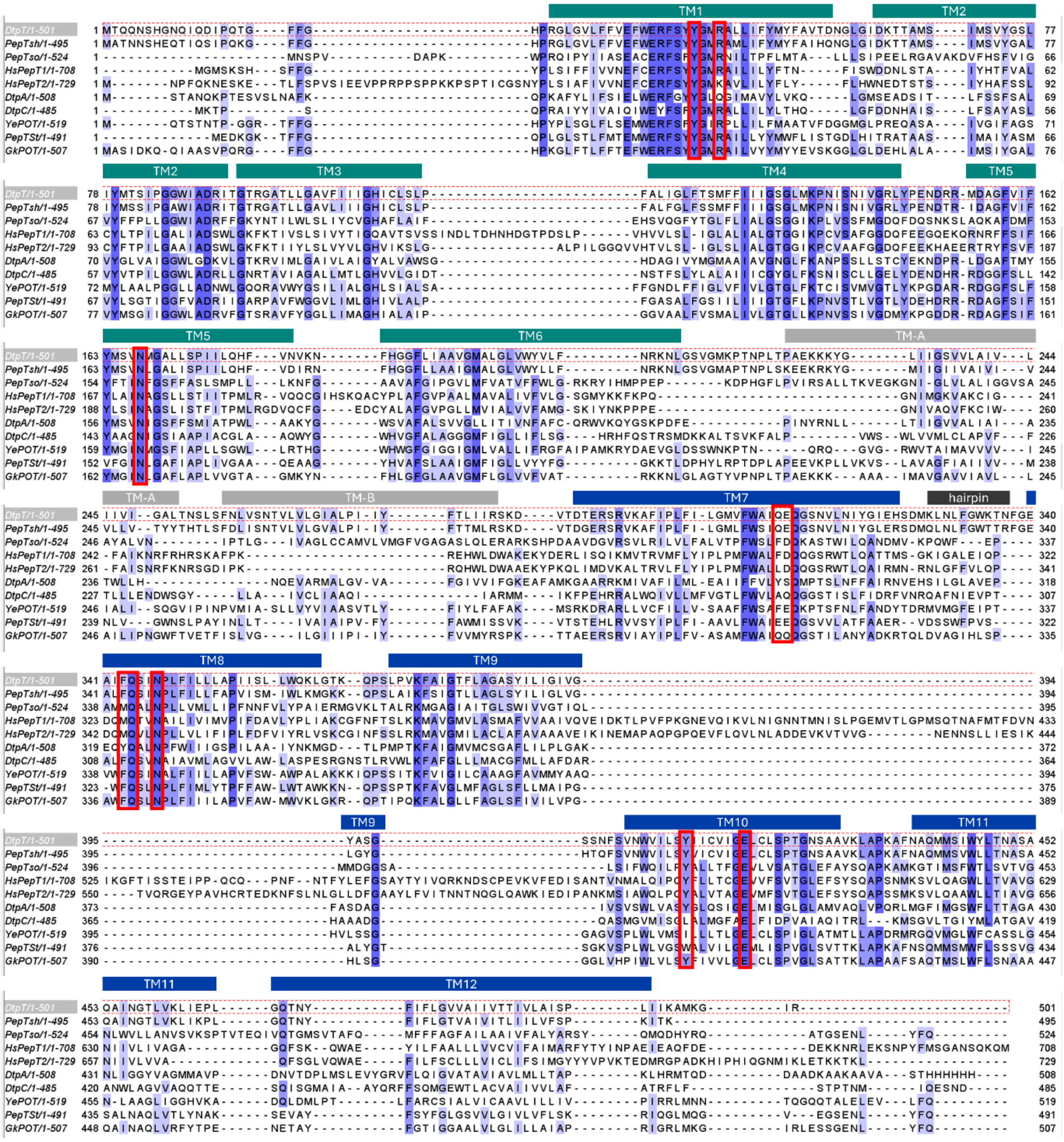

**Figure S7 – Multiple sequence alignment of DtpT and structurally characterised POTs.** Multiple protein sequence alignment was carried out by Clustal Omega (Madeira et al., 2024) and visualised in Jalview (Waterhouse et al., 2009). The sequence of DtpT (from *S. aureus* JE2) is highlighted. Sequences are given in FASTA format and coloured by identity (blue). Residues predicted to contribute toward GSH binding in DtpT are highlighted by red boxes. Regions corresponding to predicted structural elements in DtpT are also labelled above each line.

**Supplemental figure 8. Complete transport assay data for DtpT and mutant variants compared to a water-only control.**

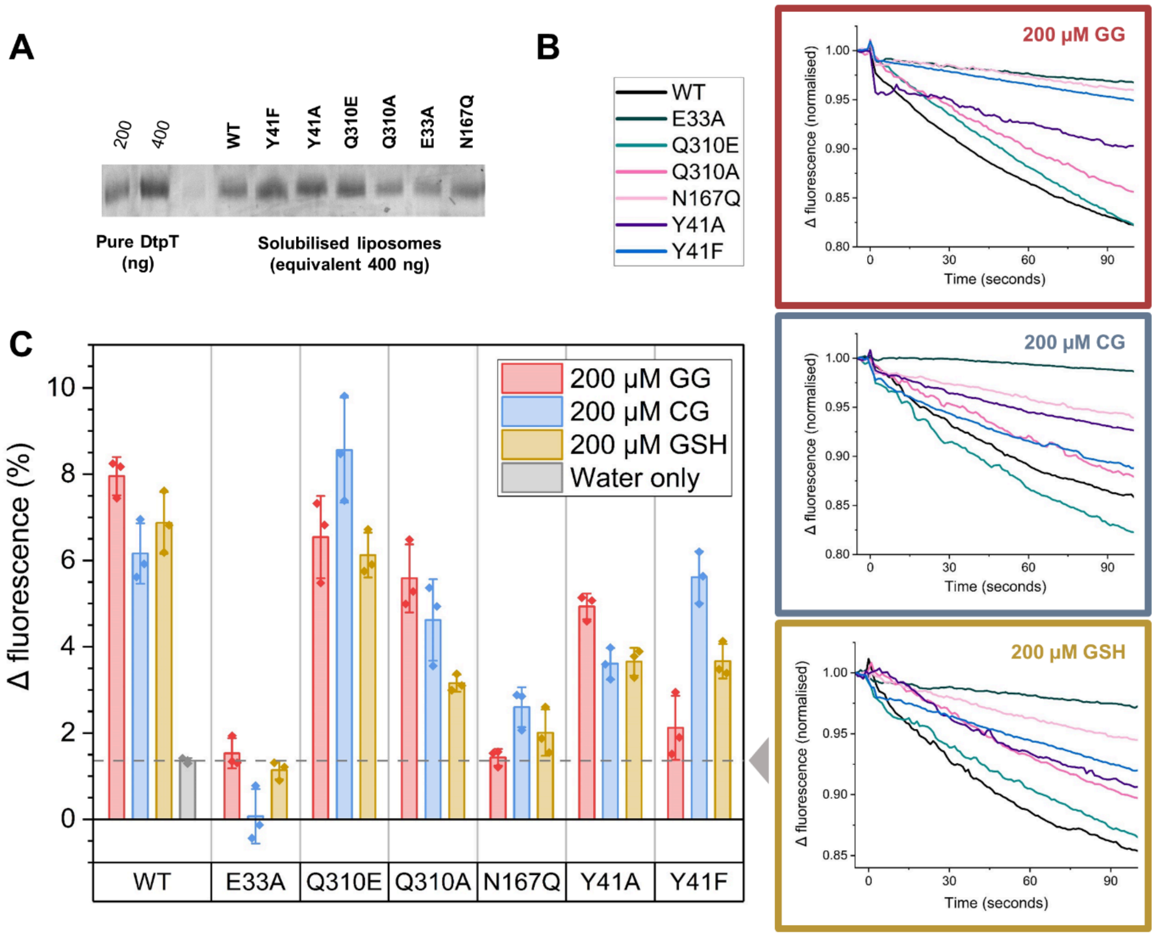

**Figure S8 – Complete transport assay data for DtpT and mutant variants compared to a water-only control.** (**A**) SDS PAGE comparison of pure DtpT against solubilised liposomes containing wild-type or mutant variants of DtpT, as labelled. Band intensities were used to normalise the final liposomal protein concentration to 0.5 mg/ml. (**B**) Complete transport assay data for each of three peptide substrates, normalised immediately before the addition of valinomycin for ease of comparison. Curves show the mean of three replicates. (**C**) Summarised transport data for DtpT and mutant variants against gly-gly, cys-gly and GSH. Transport activity is compared by quantifying the change in pyranine fluorescence over the first 30 seconds of the assay. A grey dashed line indicates the Δ fluorescence (%) recorded for DtpT liposomes in the absence of substrate (water only). Bars indicate the mean ± standard deviation

**Supplemental table 1. Strains utilised in this work**

| Strain | Organism | Background | Relevant genotype | Notes | Relevant source |
|---|---|---|---|---|---|
| **IK5** | *S. aureus* | JE2 | | | Fey et al., 2013 |
| **IK2** | *S. aureus* | JE2 | *dtpT*::$Erm^R$ | Generated via Φ85 phage transduction from NTML strain NE971 into *IK5* | Fey et al., 2013 |
| **IK163** | *S. aureus* | JE2 | *opp3A*::$Erm^R$ | Generated via Φ85 phage transduction from NTML strain NE440 into *IK5* | Fey et al., 2013 |
| **IK38** | *S. aureus* | JE2 | *dtpT::* | *ermR* casette cleared from *bursa aurealis* via allelic exchange | Bose et al., 2013 |
| **IK165** | *S. aureus* | JE2 | *dtpT*:: *opp3A*::$Erm^R$ | Generated via Φ85 phage transduction from NTML strain NE440 into *IK38* | Fey et al., 2013 |
| **GHT2288** | *E. coli* | DH5α | pSK5630 | *E. coli* / *S.aureus* shuttle vector | Grkovic et al., 2003 |
| **IK161** | *E. coli* | DH5α | $pSK56_{pT}$ | 1080 bp of sequence immediately upstream of *dtpT* in the *S. aureus* JE2 genome (incorporating the DtpT promoter sequence) was cloned into pSK5630 | This study |
| **IK170** | *E. coli* | DH5α | $pSK56_{pT}$*dtpT* | The entire *dtpT* open reading frame and 1080 bp of upstream sequence from *S. aureus* JE2 was cloned into pSK5630 | This study |
| **IK12** | *S. aureus* | RN4220 | | Cloning intermediate strain | Kreiswirth et al., 1983 |
| **IK176** | *S. aureus* | JE2 | $pSK56_{pT}$ | | This study |

| | | | | | |
|---|---|---|---|---|---|
| **IK177** | *S. aureus* | JE2 | *dtpT::* + pSK56$_{pT}$ | | This study |
| **IK204** | | | *dtpT::* + pSK56$_{pT}$*dtpT* | | This study |
| **IK209** | *S. aureus* | JE2 | *dtpT*:: *opp3A*:: Erm$^{R}$ + pSK56$_{pT}$ | | This study |
| **IK212** | | | *dtpT*:: *opp3A*::Erm$^{R}$ + pSK56$_{pT}$*dtpT* | | This study |
| **IK208** | *S. aureus* | JE2 | *gisD*::Erm$^{R}$ | Generated via Φ85 phage transduction from NTML strain NE215 into *IK5* | Fey et al., 2013 |
| **IK207** | | | dtpT:: *gisD*::Erm$^{R}$ | Generated via Φ85 phage transduction from NTML strain NE215 into *IK38* | Fey et al., 2013 |
| **IK213** | *S. aureus* | JE2 | *dtpT*:: *gisD*:: Erm$^{R}$ + pSK56$_{pT}$ | | This study |
| **IK214** | | | *dtpT*:: *gisD*::Erm$^{R}$ + pSK56$_{pT}$*dtpT* | | This study |
| **IK206** | *S. aureus* | JE2 | SAUSA300_0060 ::Erm$^{R}$ | Generated via Φ85 phage transduction from NTML strain NE806 into *IK5* | Fey et al., 2013 |
| **IK32** | *E. coli* | BL21(DE3) | pWaldo | *E. coli* membrane protein expression vector containing C-terminal octa-His GFP-tag | Drew et al., 2006 |
| **IK81** | *E. coli* | C43 | | | Miroux and Walker, 1996 |
| **IK98** | *E. coli* | C43 | pWaldo-DtpT | | This study |
| **IK187** | | | pWaldo-DtpT$_{Y41F}$ | | This study |
| **IK188** | | C43 | pWaldo-DtpT$_{Y41A}$ | | This study |

| | | | | | |
|---|---|---|---|---|---|
| **IK191** | | | pWaldo-DtpT$_{Q310E}$ | | This study |
| **IK197** | | | pWaldo-DtpT$_{Q310A}$ | | This study |
| **IK189** | | | pWaldo-DtpT$_{E33A}$ | | This study |
| **IK190** | | | pWaldo-DtpT$_{N167Q}$ | | This study |

**Supplemental table 2. Primers utilised in this work**

| Primer number | Target | Sequence | Function |
|---|---|---|---|
| 1 | *dtpT* flanking regions | GGGCAAGTTCACACCACG | diagnostic PCR |
| 2 | | GTTAACCGTGCCATCACCG | |
| 3 | *opp3A* flanking regions | CCTTAGCTCCTGAAAATCTCGTCG | diagnostic PCR |
| 4 | | GCGGATTTGAACGTTGCGAAG | |
| 5 | *gipD* flanking regions | GGTGTTTGTATTATGTTCGTCG | diagnostic PCR |
| 6 | | TGGTTCTAACACATTCAATGCC | |
| 7 | *Bursa aurealis* Tn | GCTTTTTCTAAATGTTTTTTAAGTAAATCAAGTAC | diagnostic PCR |
| 8 | pWaldo | GAAGCAGCCCAGTAGTAGG | diagnostic PCR/sequencing |
| 9 | | GTGTTGGCCATGGAACAGG | |
| 10 | pSK5630 | CCTTAGCTCCTGAAAATCTCGTCG | diagnostic PCR/sequencing |
| 11 | | GCGGATTTGAACGTTGCGAAG | |
| 14 | pWaldo cloning site | GTCGAGTCTCCTTCTTAAAGTTAAAC | vector linearisation |
| 15 | | GAAAACCTGTACTTCCAGGGTCA | |
| 16 | pSK5630 cloning site | GATCCGTCGACCTGCAGC | vector linearisation |
| 17 | | CCCTGGCAGTTTATGGCGG | |
| 20 | *dtpT* | (TAACTTTAAGAAGGAGACTCG)ACATGACACAACAAAACTCCCA | Cloning into pWaldo |
| 21 | | (CCTGGAAGTACAGGTTTTC)ACGTATACCTTTCATCGCTTTGA | |
| 22 | *dtpT* | (CCGCCATAAACTGCCAGGG)ATTAACGTATACCTTTCATCGC | cloning into pSK5630 |
| 23 | *dtpT* promoter region | (GCTGCAGGTCGACGGATC)GGGTCAAATTGGTGTTACTT | |
| 24 | | (CCGCCATAAACTGCCAGGG)CATGTATACATCCCATCCTTTC | |
| 27 | *dtpT* | CGTACCAATTGCAAATTTTACTGG | diagnostic PCR |
| 28 | *dtpT* | CTGGGCTATTgcaGAACAAGGGTC | Q310A mutation |
| 29 | | CTGGGCTATTgaaGAACAAGGGTC | Q310E mutation |
| 30 | | AACACCATTCCAAGAATAAATAATG | Q310 reverse |
| 31 | | GTTTAGTTATtttGGCATGCGTG | Y41F mutation |
| 32 | | GTTTAGTTATgctGGCATGCGTGC | Y41A mutation |
| 33 | | CTTTCCCAGAACTCTACAAAG | Y41 reverse |
| 34 | | CTTCTTTGTAgcgTTCTGGGAAAG | E33A mutation |
| 35 | | AGTACGCCTAGTCCTC | E33 reverse |
| 36 | | TATGTCAGTTcaaATGGGTGCATTATTATC | N167Q mutation |
| 37 | | TAGAAAATAACAAAACCTGC | N167 reverse |

**Supplemental table 3. Composition of CDMG (adapted from Hussain et al., 1991)**

| Ingredients | mg/L |
| --- | --- |
| **Group 1** | **Adjust pH to 7.2, then autoclave in 700 ml** |
| $Na_2HPO_4$ | 8000 |
| $KH_2PO_4$ | 3000 |
| l-Aspartic acid | 150 |
| l-Alanine | 100 |
| l-Arginine | 150 |
| l-Cysteine | 75 |
| Glycine | 100 |
| l-Glutamic acid | 150 |
| l-Histidine | 100 |
| l-Isoleucine | 150 |
| l-Lysine | 100 |
| l-Leucine | 150 |
| l-Methionine | 100 |
| l-Phenylalanine | 100 |
| l-Proline | 150 |
| l-Serine | 100 |
| l-Threonine | 150 |
| l-Tryptophan | 100 |
| l-Tyrosine | 100 |
| l-Valine | 150 |

| Ingredients | mg/L |
| --- | --- |
| **Group 2** | **Autoclave in 280 ml** |
| d-Glucose | 2 000 |
| $MgSO_4 \cdot 7H_2O$ | 500 |

| | |
| --- | --- |
| **Group 3** | |
| **Filter sterile to 100x stock** | **+ 10 ml** |
| Biotin | 0.1 |
| Nicotinic acid | 2 |
| D-Pantothenic acid, Ca salt | 2 |
| Pyridoxal | 4 |
| Thiamine hydrochloride | 2 |
| Pyridoxamine dihydrochloride | 4 |
| **Group 4** | |
| ***Filter sterile to 100x stock*** | **+ 10 ml** |
| $CaCl_2 \cdot 6H_2O$ | 10 |
| $MnSO_4$ | 5 |
| $(NH_4)_2SO_4 \cdot FeSO_4 \cdot 6H_2O$ | 6 |

**SM1 – Supplemental methods supporting figure S8**

**Ethics Statement**

This study was conducted in accordance with the recommendations in the Guide for the Care and Use of Laboratory Animals of the National Institutes of Health. The approved protocol, PROTO202200474, was reviewed by the Animal Care and Use Committee at Michigan State University.

**Strains used in animal experiments**

The Δ*gis* and *gisB*::Tn were previously described in Lensmire et al., 2023. The *dtpT*::Tn and Δ*gis dtpT*::Tn strains were generated via phage transduction into the WT JE2 strain background or the previously published Δ*gis* strain background. All strains used in animal experiments are described in the following table:

| Organism | Background | Relevant genotype | Notes | Relevant source |
|---|---|---|---|---|
| *S. aureus* | JE2 | | | Fey et al., 2013 |
| | | *dtpT*::Tn | Generated via phage transduction from NTML strain NE971 into JE2. Functionally identical to IK2 but generated independently by NDH. | Fey et al., 2013 |
| | | *gisB::*Tn | Generated via phage transduction from NTML strain NE215 into JE2. | Fey et al., 2013 |
| | | *Δgis* | | Lensmire et al., 2023 |
| | | *Δgis dtpT::Tn* | Generated via phage transduction from NTML strain NE440 into *Δgis* | Lensmire et al., 2023 |

**Single strain murine systemic infections**

WT, *dtpT*::Tn, or Δ*gis dtpT*::Tn mutant strains were cultured in TSB overnight at 37°C, diluted 1:100 into TSB, and cultured for 3 h at 37°C at 225 rpm shaking. Cultures were pelleted, washed with PBS, and normalized to $OD_{600}$ equal to 0.4. Thirty female 8-week-old BALB/cJ mice were retro-orbitally infected with $10^7$ CFUs and the infection proceeded for

96 h after which heart, liver, and kidneys were collected and homogenized in 1 mL PBS. Organ homogenates were serially diluted and plated onto TSA. Bacterial burdens were quantified as CFUs $mL^{-1}$. Infections were performed at Michigan State University under the principles and guidelines described in the Guide for the Care and Use of Laboratory Animals. Animal work was followed as approved by Michigan State University Institutional Animal Care and Use Committee (IACUC) approved protocol number PROTO202200474.

**Murine systemic competition infections**

WT, *dtpT*::Tn, *gisB*:Tn and Δ*gis dtpT*::Tn mutant strains were grown in TSB overnight at 37°C, subcultured 1:100, and grown in TSB for 3 h at 37°C and 225 rpm. Strains were washed in PBS and normalized to an $OD_{600}$ equal to 0.4. The competitions were prepared by mixing equal volumes of WT with the *dtpT*::Tn, *gisB*:Tn or Δ*gis dtpT*::Tn mutant strains. The input ratio was quantified by serially diluted the mixture and plating onto TSA and TSA supplemented with 10 μg $mL^{-1}$ erythromycin ($erm^{10}$) to discern between WT and the mutant strain. The transposon-harboring mutant strains utilized in these experiments harbor an erm resistance cassette within the transposon. WT CFU were calculated by subtracting CFU generated on TSA-$erm^{10}$ from CFU produced on TSA. Fifteen female 8-week-old BALB/cJ mice were retro-orbitally infected with 100 μL containing the mixture of $10^7$ CFU of WT and the indicated mutant strain. After 96 h, the heart, liver, and kidneys were collected and homogenized in 1 mL PBS. Homogenates were serially diluted and plated onto TSA or TSA-$erm^{10}$. Competitive indices were calculated as dividing the WT:mutant strain output CFU ratio by the WT:mutant input CFU ratio.

**Statistical Analysis**

All data were analyzed using GraphPad Prism v9.1.1.
