## Supplementary figures and images for "Defined roles for the *Staphylococcus aureus* POT transporter DtpT in di/tripeptide uptake and glutathione utilisation inside human macrophages"

### Supplemental Data SD1

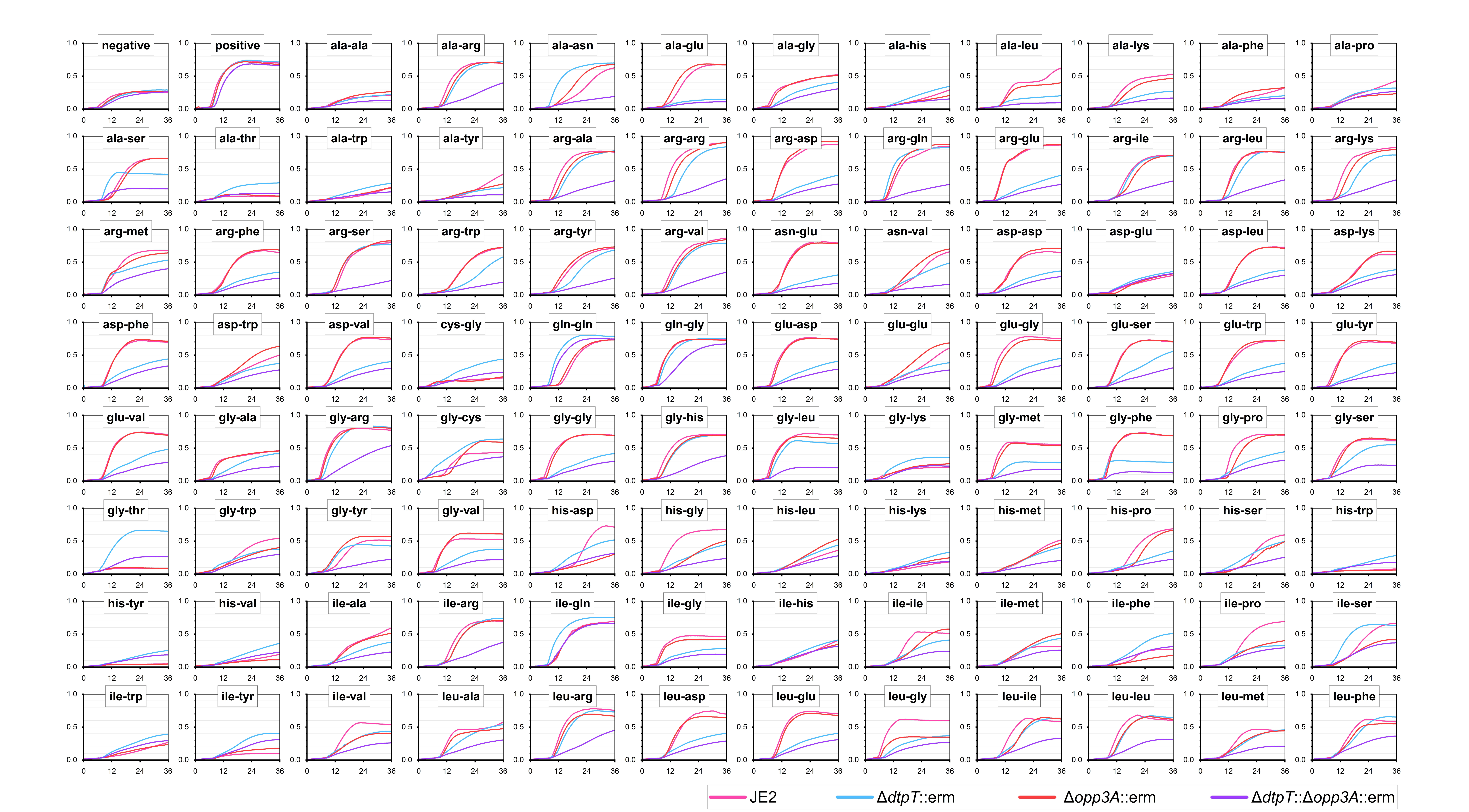

### Supplemental Data SD2

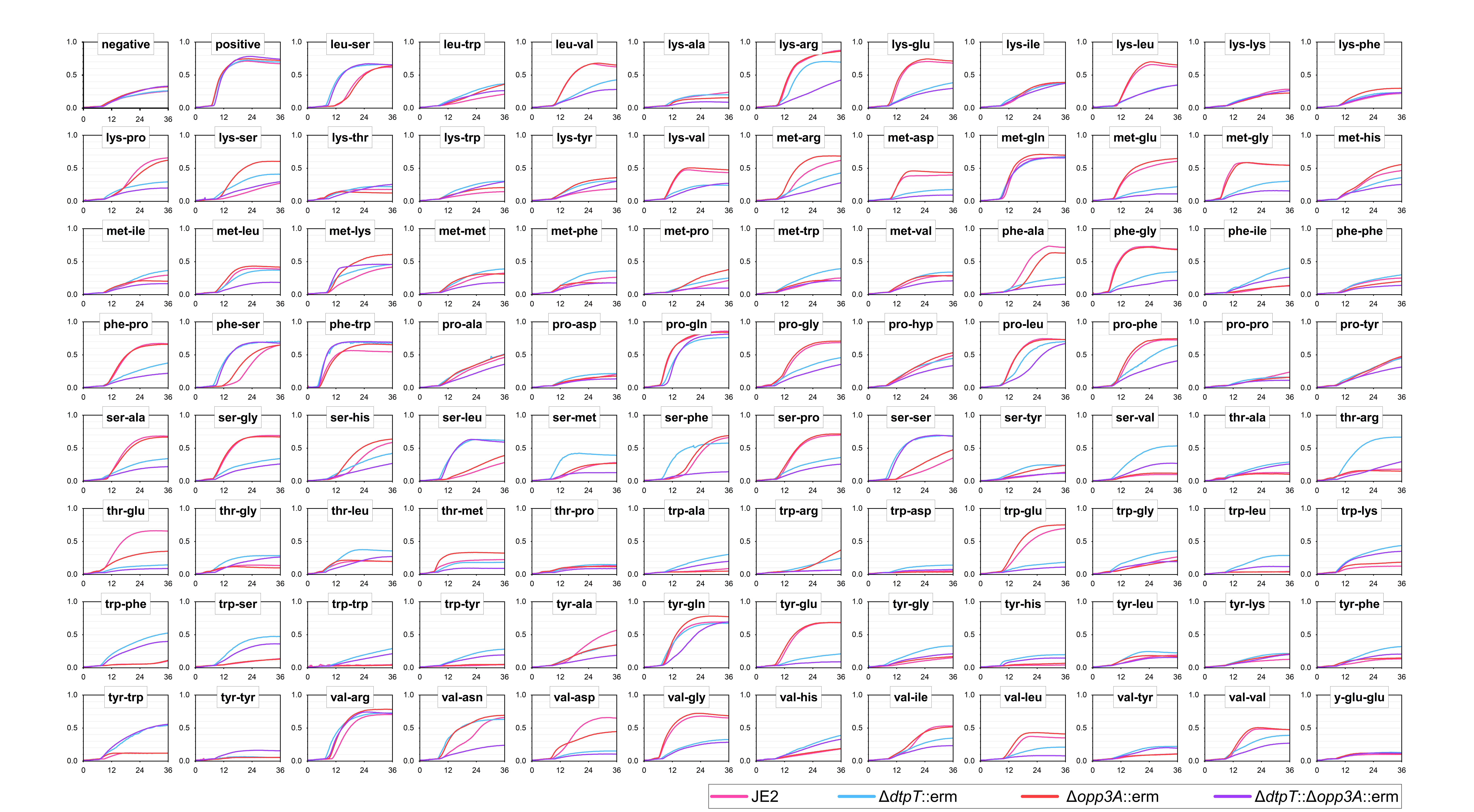

### Supplemental Data SD3

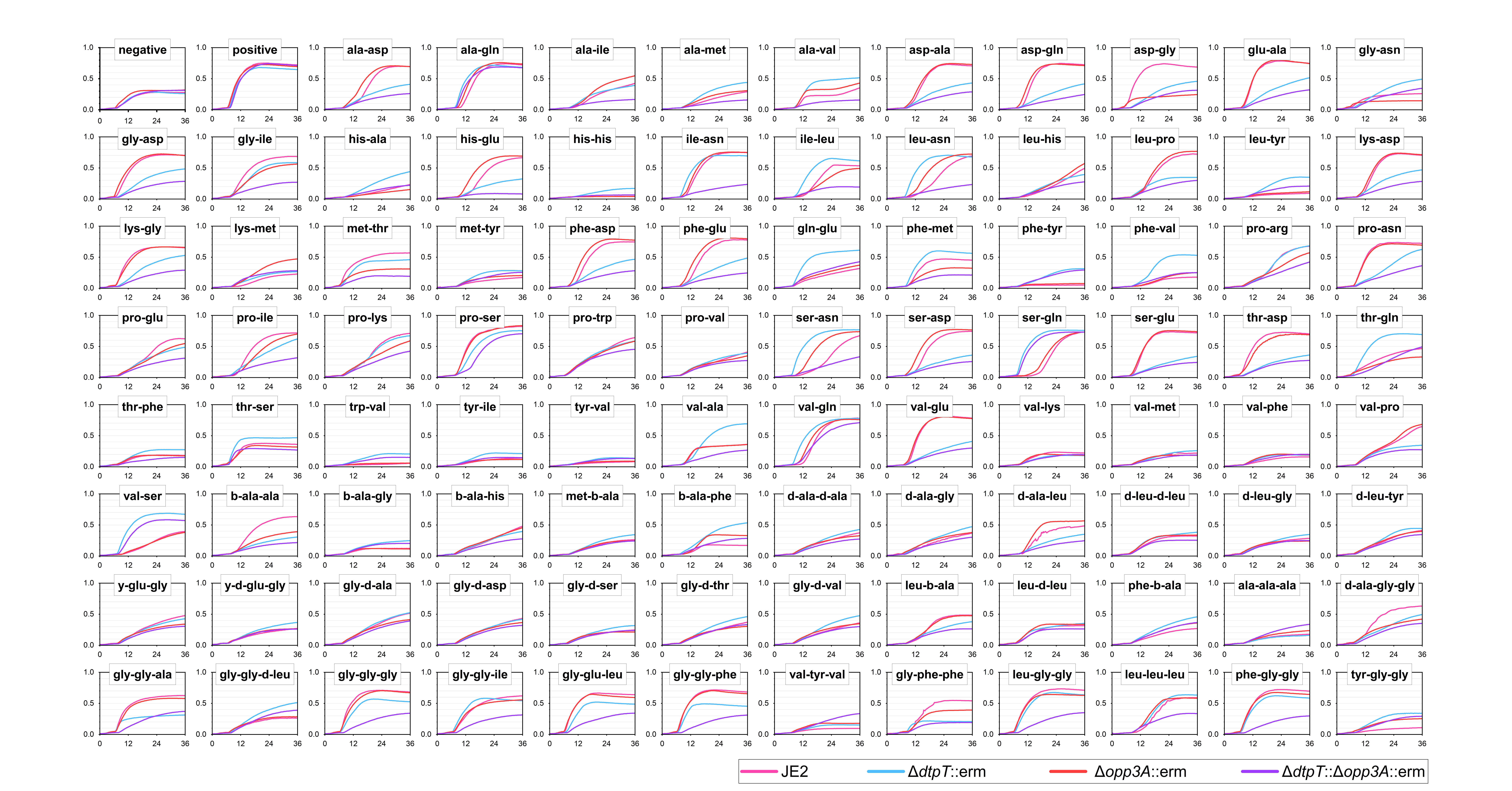
